# Kinetic co-evolutionary models predict the temporal emergence of HIV resistance mutations under drug selection pressure

**DOI:** 10.1101/2022.11.30.518575

**Authors:** Avik Biswas, Indrani Choudhuri, Eddy Arnold, Dmitry Lyumkis, Allan Haldane, Ronald M. Levy

## Abstract

Drug resistance in human immunodeficiency virus (HIV) is a pervasive problem that affects the lives of millions of people worldwide. Although records of drug-resistant mutations (DRMs) have been extensively tabulated within public repositories, our understanding of the evolutionary kinetics of DRMs and how they evolve together remains limited. Epistasis, the interactions between a DRM and other residues in HIV protein sequences, is found to be key to the temporal evolution of drug resistance. We use a Potts sequence-covariation statistical-energy model of HIV protein fitness under drug selection pressure, which captures epistatic interactions between all positions, combined with kinetic Monte-Carlo simulations of sequence evolutionary trajectories, to explore the acquisition of DRMs as they arise in an ensemble of drug-naïve patient protein sequences. We follow the time course of 52 DRMs in the enzymes protease, reverse transcriptase, and integrase, the primary targets of antiretroviral therapy (ART). The rates at which DRMs emerge are highly correlated with their observed acquisition rates reported in the literature when drug pressure is applied. This result highlights the central role of epistasis in determining the kinetics governing DRM emergence. Whereas rapidly acquired DRMs begin to accumulate as soon as drug pressure is applied, slowly acquired DRMs are contingent on accessory mutations that appear only after prolonged drug pressure. We provide a foundation for using computational methods to determine the temporal evolution of drug resistance using Potts statistical potentials, which can be used to gain mechanistic insights into drug resistance pathways in HIV and other infectious agents.

**Significance:** HIV affects the lives of millions of patients worldwide; cases of pan-resistant HIV are emerging. We use kinetic Monte-Carlo methods to simulate the evolution of drug resistance based on HIV patient-derived sequence data available on public databases. Our simulations capture the timeline for the evolution of DRMs reported in the literature across the major drug-target enzymes – PR, RT, and IN. The network of epistatic interactions with the primary DRMs determines the rate at which DRMs are acquired. The timeline is not explained by the overall fitness of the DRMs or features of the genetic code. This work provides a framework for the development of computational methods that forecast the time course over which drug resistance to antivirals develops in patients.

## Introduction

The human immunodeficiency virus type 1 (HIV-1) currently infects ∼40 million people worldwide. In the absence of a cure, antiretroviral therapy (ART) presents the primary treatment option (1). However, all antiretroviral drugs, including those from newer drug classes, are at risk of becoming partially or fully inactive due to the emergence of drug-resistant mutations (DRMs) (2-5). The rapid mutation rate of HIV plays a major role in the failure of ARTs amongst infected patients, leading to DRMs occurring in response to drug selection pressure (6, 7).

The viral enzymes protease (PR), reverse transcriptase (RT), and Integrase (IN), which are encoded by the *Pol gene* of HIV-1, have been the major focus of ART over the past several decades (8-14). The fitness landscape of these enzymes is determined by the combined effects of the host immune response and selection pressure from ART, and how these interplay with the proteins’ structure, function, thermodynamics, and kinetics (15-20). As a result, complex mutation profiles often arise in these proteins located both near and distal from the active site (17, 21, 22). These profiles can be observed in HIV patient protein sequences that are available on large public databases such as the Stanford HIV drug resistance database (HIVDB) (23) and the Los Alamos HIVDB (24, 25), from which we can derive specific patterns and relationships.

Primary DRMs generally occur with a fitness penalty in viral sequences found in drug naïve patients. The effect of a mutation, however, is dependent on the entire genetic background in which it occurs, a phenomenon known as ‘epistasis’. Due to epistatic interactions with the sequence background, primary DRMs can become very favorable (26, 27) in sequence backgrounds in which accessory mutations accumulate, such that there is a fitness penalty for reversion to the wild-type residue that leads to evolutionary trapping or “entrenchment” of the primary mutation (28-32). There is also feedback between the appearance of primary and background mutations, with the accumulation of accessory mutations increasing the likelihood of the primary mutation to arise (“contingency”), and vice versa. In the presence of drug pressure, this leads to a complex interplay between the functions of the primary and accessory mutations (33, 34). Theoretical considerations suggest that epistasis can slow the rate of evolution by creating a rugged fitness landscape with low-fitness intermediate states forming barriers between local fitness optima (35, 36). Here, we study such phenomena using empirical data.

Studies have illustrated the effects of epistasis on the fitness landscape of HIV proteins (29, 34, 37-40), but it is unclear why some DRMs are acquired rapidly while others are acquired much more slowly, and how this is influenced by the epistatic network. Recently, we introduced a kinetic model that describes the evolution of HIV sequences on a fitness landscape constructed using a Potts sequence-covariation statistical energy model (41) inferred from drug experienced patient protein sequences in the Stanford HIVDB (7, 23). This kinetic evolution model has the key feature that it models epistasis between all positions in the sequences. We have previously shown that this feature is critical for making the model numerically consistent with the observed between-host sequence mutation co-variation statistics obtained from public repositories, such as the Stanford HIVDB. When simulating many long evolutionary trajectories in parallel by our method starting from a drug-naïve sequence ensemble and collecting the final sequences, the mutational statistics (frequencies of single point and higher-order combinations of mutations) of the generated sequences match those of the drug-experienced dataset that was used to train the fitness model. To establish a baseline for understanding how epistasis affects kinetics associated with the development of drug resistance, we previously used this model to follow the kinetics of a DRM within the drug-experienced ensemble of patient protein sequences and concluded that epistasis has a strong effect on evolutionary dynamics (37). However, the emergence of drug resistance is best understood in the context of a changing environment as the virus is newly exposed to drug treatment. In the current work, we focus on the kinetics of the emergence of drug-resistance mutations in an ensemble of drug-naïve HIV patient sequences evolving under the influence of newly applied drug pressure.

Our goal is to use the kinetic model to probe the relative times at which DRMs arise in HIV under drug selection pressure. We focus on 52 DRMs in the three HIV drug-target enzymes coded by the *Pol* gene (23). These 52 DRMs are chosen based on two primary criteria: they are classified in the Stanford HIVDB as ‘primary’ mutations that generally affect drug susceptibility ≥5-10 fold on their own (23), and they are observed in drug-experienced patients with frequencies ≥1%. The principal result of our study is that the rates at which DRMs are acquired in initially drug-naïve strains predicted using our model are highly correlated with the corresponding observed acquisition rates that are reported in the literature. This suggests that the rates at which DRMs are acquired depend strongly on epistatic interactions between the focal mutation and the other residues in the sequence background. The acquisition rates cannot be explained by the equilibrium frequency of a DRM, which is a proxy for its fitness in the drug-experienced patient population after averaging over sequence backgrounds or by features of the genetic code such as nucleotide changes (Δnuc) at the codon level, or transitions (Ti) vs. transversions (Tv). We propose that some DRMs are acquired more slowly because they face an “epistatic barrier”, and outline how this arises. The Potts model parameterized on drug-experienced HIV patients combined with kinetic Monte Carlo techniques is a powerful predictor of the relative DRM acquisition times leading to drug resistance.

## Results

### Relative rates at which HIV DRMs are acquired with a kinetic Monte-Carlo evolution model match the literature

We follow the temporal evolution of a set of primary DRMs in HIV-1 protease (PR), reverse transcriptase (RT), and integrase (IN) using a kinetic Monte-Carlo method to evolve, in parallel, an ensemble of initially drug-naïve consensus sequences. Our kinetic Potts model employs a coarse-grained representation of the intra-host HIV evolutionary process, as described in more detail in Computational Methods. During HIV infection, many viral variants are present in a host, but the sequences of these within-host variants are generally ∼99% identical (42). This stands in contrast to the sequence identity between evolving viral populations in different hosts (between-hosts), which can be much lower (e.g. 90% for PR) (43). These observations suggest that a host’s viral population can be represented at coarse grain by its population consensus sequence. Indeed, in many HIV sequence datasets, such as derived from the Stanford HIVDB, effectively a population consensus sequence is sampled from each host by averaging over the viral diversity within the host. We model the evolution of an ensemble of consensus sequences representing multiple host populations, approximating this process as a series of point-mutation events occurring at a constant rate in each consensus sequence, consistent with observations (44), and the mutations are either fixed or lost according to the fitness landscape inferred based on between-host sequence data. While this model coarse-grains several details of the intra-host HIV evolutionary dynamics, for instance clonal competition, recombination, immune-pathogen co-evolution, spatial and temporal drug heterogeneity, and nucleotide-level biases, it faithfully reproduces the observed pairwise and higher-order mutation patterns of the data.

We evolve the ensemble of sequences over a fitness landscape represented by a drug-experienced Potts statistical energy model that captures the sequence covariation due to selection pressure in the presence of drugs. The Potts ‘statistical’ energy (E(S)) of a sequence S (Computational Methods section) predicts how sequences will appear in the dataset with probability *P*(*S*) ∝ *e*^−*E*(*S*)^ such that sequences with favorable statistical energies are more prevalent in the multiple sequence alignment, and thus more fit. This overall fitness predicted by the Potts model, which is inferred based on observed mutation prevalence, is the net result of many phenotypic contributions including both the “replicative fitness” and “transmission fitness” commonly measured by virological assays (45-47). A key feature of the Potts model of fitness is that the effect of a mutation on E(S) is ‘background-dependent’: a mutation at one position will affect mutations at all other positions both directly and indirectly through chains of epistatic interactions involving one or many intermediate residues.

Figure 1 illustrates how drug-exposure drives the appearance of DRMs in general, as demonstrated with the enzyme RT. The distribution of the total number of mutations per sequence measured with respect to the wild-type (WT) HIV-1 subtype-B consensus sequence (hereafter referred to more simply as the WT) evolves from a narrow initial distribution in drug-naïve sequences with a peak at ∼6 mutations and a maximum of ∼20 mutations to a much broader distribution in drug-experienced sequences with a peak at ∼12 mutations and a maximum of more than 30 mutations. Corresponding plots for PR and IN are shown in **Fig. S1** and **S2** (**SI**). We note that IN is more conserved under drug pressure than either PR or RT.

**Figure 1.**
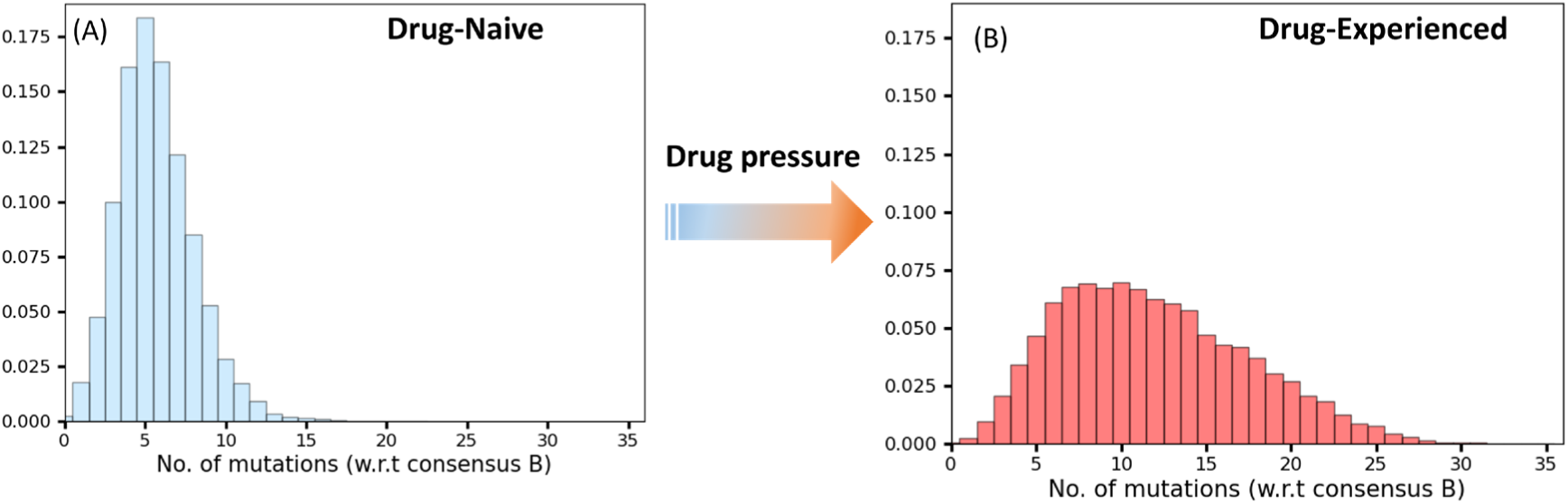
The distribution of the total number of mutations in HIV-1 RT in (A) the drug-naïve multiple sequence alignment (MSA), and (B) the drug-experienced MSA available from the Stanford HIVDB (22, 24).

We compute a characteristic acquisition time for each primary DRM in PR, RT, and IN from the kinetic Monte-Carlo (KMC) simulations using an exponential fit for the change in the frequency of the DRM starting from the initial drug-naïve ensemble and ending in the final drug-experienced state (**Fig. S3**, **SI**). We follow 52 primary DRMs with a wide range of acquisition times in the clinical literature: 13 protease inhibitor (PI) DRMs, 14 nucleoside-analog RT inhibitor (NRTI) DRMs, 11 non-nucleoside RT inhibitor (NNRTI) DRMs, and 14 integrase strand-transfer inhibitor (INSTI) DRMs. Fig. 2 shows the correspondence between the acquisition times to acquire these 52 major DRMs estimated from the KMC simulations, alongside the corresponding timelines reported in the literature (**Fig. S3**, **Table S1-S4**, **SI**). The Spearman correlation coefficients between the KMC simulated and observed DRM acquisition times are listed in **Table 1**, and correlation plots are shown in **Fig. S4** (**SI**). The average Spearman correlation coefficient for all 52 DRMs in the three HIV viral enzymes is ρ=0.85, p<<0.001 (for individual enzymes, ρ=0.75, 0.92, and 0.90 for PR, RT, and IN, respectively). The very strong correlation between the predicted and observed acquisition times for primary DRMs is noteworthy. The acquisition times for each DRM in PR, RT (NRTI-selected), RT (NNRTI-selected), and IN are shown in **Table S1-S4 SI)**. The time span to acquire DRMs is large, and the fastest DRMs are acquired ∼20 times more rapidly than the slowest DRMs. To illustrate the temporal evolution of DRMs with a contrasting timeline of resistance, we divide the primary DRMs into three categories based on their acquisition times in the literature: fast, between 0-3 months; intermediate, between 4-5 months; and slow, more than 6 months. The acquisition times (*τ*) from our simulations are correspondingly classified for each category, between ∼1-10 for fast, ∼10-24 for intermediate, and >24 for slow.

**Figure 2:**
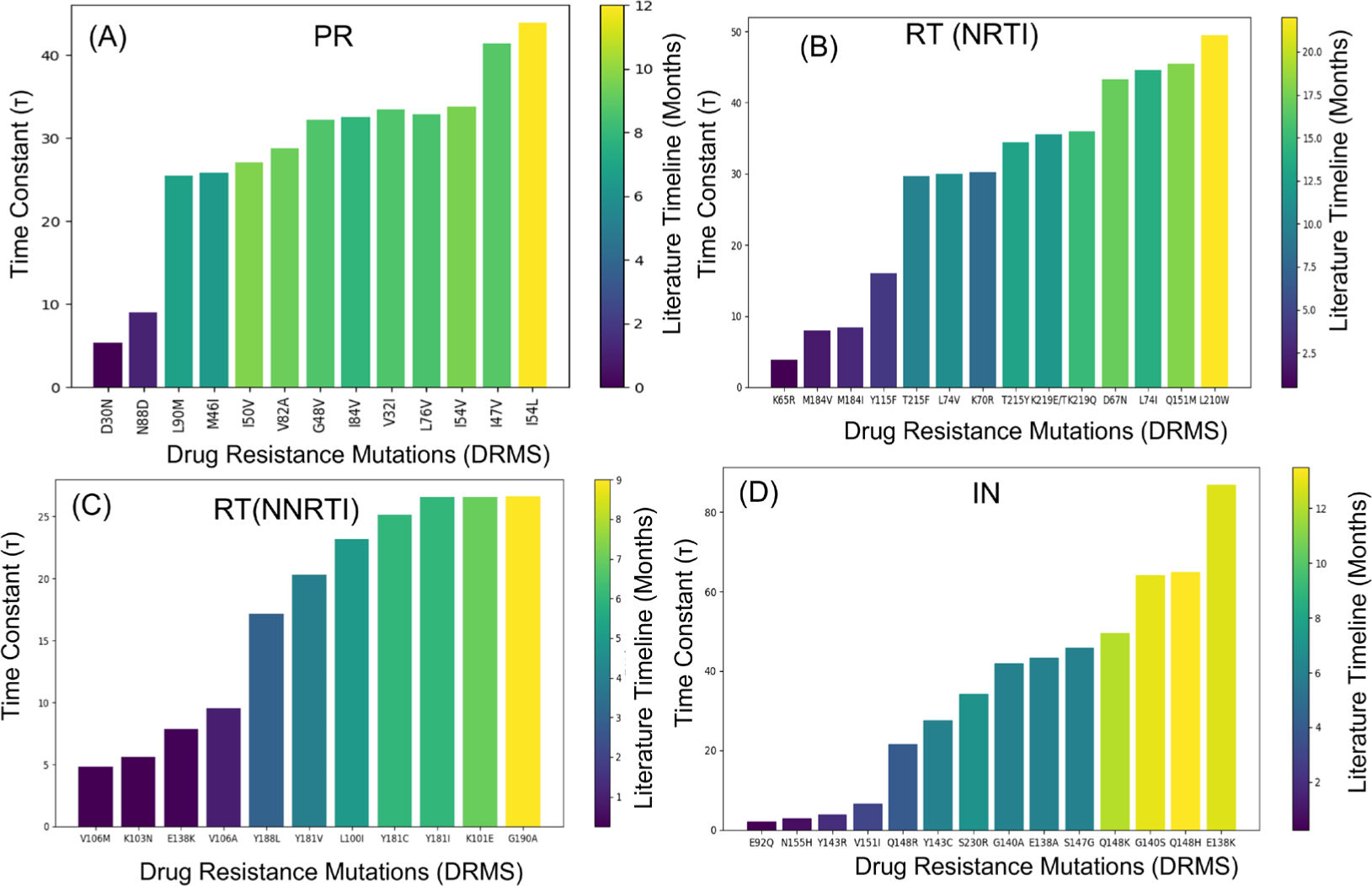
The literature correspondence with the KMC acquisition time of the major (A) PR DRMs in response to PIs, (B) RT DRMs in response to NRTI, (C) RT DRMs in response to NNRTI, and (D) IN DRMs in response to INSTIs. The height of each bar represents the time constant (τ) calculated from KMC simulations. The color bar represents the time required for each DRM to emerge in the patient’s population under drug selection pressure, which is collected from the literature.

**Table 1:** Spearman’s rank correlation coefficient (ρ) test.

| Models | Targets | $\rho$ | p-value |
| --- | --- | --- | --- |
| KMC Predicted Time constant ($\tau$) | All DRMs | 0.85 | $1 \times 10^{-22}$ |
| | PR | 0.75 | $1 \times 10^{-05}$ |
| | RT | 0.92 | $1 \times 10^{-16}$ |
| | IN | 0.90 | $1 \times 10^{-08}$ |
| Mutant Prevalence (Drug Experienced Frequency) | All DRMs | 0.26 | 0.05 |
|  | PR | 0.30 | 0.29 |
|  | RT | 0.47 | 0.02 |
|  | IN | 0.02 | 0.93 |
| Genetic Code (Ti vs. Tv) | All DRMs | 0.13 | 0.33 |
|  | PR | 0.18 | 0.54 |
|  | RT | 0.23 | 0.25 |

Among the NRTI selected primary DRMs, 3 are acquired rapidly, while 10 are acquired slowly. The DRMs K65R, M184V, and M184I are acquired within ∼1.5-3 months after initiation of therapy with KMC time constants ranging between ∼4-8 KMC time units, whereas T215F/Y, L74V/I, K70R, K219E/Q, D67N and L210W are acquired between 9-20 months after initiation of therapy with KMC time constants between 30-50 KMC time units (**Table S2-S3, SI**). The relative times required for acquisition of the major drug resistance mutations in PR and IN, (**Table S1** and **Table S4**, **SI**) are also recapitulated by the KMC simulations. We conclude from the results summarized in (**Fig. S3** and **Table S1-S4**, **SI**), that the KMC model of evolution of drug resistance mutations in RT captures fundamental kinetic features of the mutational landscape for this enzyme evolving under drug pressure, which distinguishes DRMs that are acquired slowly from those acquired rapidly, as reported in the literature.

The KMC simulations correspond to a fitness based coarse-grained epistatic model of the HIV evolutionary process under drug selection pressure. We have found that despite this coarse-graining, the model is an excellent predictor of DRM acquisition times, although it is still possible this coarse-grained model leaves out some contributors to the acquisition time. In the Computational Methods, we present tests carried out to determine if acquisition times are affected by other features that have been suggested to affect these times but are not explicitly accounted for in our model, including biases due to kinetic effects associated with nucleotide transition vs transversion rates and differences in the rates of codon changes due to single vs double-nucleotide mutations. From the low correlation between these properties and the times to acquire corresponding DRMs reported in the literature (**Table 1**), we conclude that these are not major determinants of the rates at which drug resistance is acquired. Our results, **Table 1** and Fig. 2 (**Table S1**-**S4**, **SI**) show that the epistatic coarse-grained kinetic model is successfully able to capture the kinetic behavior of a wide range of primary DRMs in different HIV drug-target proteins (PR, RT and IN). The key determinant of the rate at which DRMs are acquired is epistasis caused by the structural and functional constraints on HIV proteins, which induce sequence co-variation that the KMC model accounts for explicitly.

### Slowly acquired DRMs have an epistatic barrier to resistance

We will show in this section that the DRMs that are acquired slowly are contingent on the acquisition of accessory mutations that must arise first as the drug naïve ensemble evolves under newly applied drug selection pressure, while DRMs that are acquired rapidly have higher fitness after drug pressure is applied even before accessory mutations accumulate. Regardless of whether the DRMs are acquired rapidly or slowly, accessory mutations are eventually acquired, and they function to trap or entrench the DRMs (**Fig. S5**) in a subset of the drug-experienced sequence ensemble where they appear with high probability (34).

We explore the role of epistasis in the evolution of drug resistance starting from the drug-naïve ensemble by defining an “adaptive-frequency”, 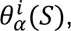 which reflects the likelihood of a mutation α to occur at position *i* in a specific sequence *S*, and differs for each sequence. In other words, if residue positions other than *i* are held fixed in sequence *S*, the adaptive frequency 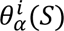 measures the likelihood of position *i* mutating to residue α relative to other possible residues at that position in the drug-experienced fitness environment. It can be thought of as a proxy for the viral fitness of the DRM in that sequence background under drug selection pressure. Using the Potts epistatic model, we define the adaptive frequency for a mutation *α* at position *i* in a sequence *S* as:

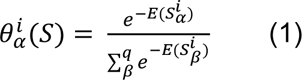

where 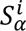 is the sequence *S* with position *i* mutated to character *α, E(S)* is the Potts statistical energy of sequence *S* parameterized on the drug-experienced patient sequence data, which includes both position-dependent ‘field’ and epistatic ‘coupling’ terms between the focal position and all other positions; the lower sum runs over all possible residues at position *i*. 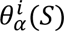 can be interpreted as the equilibrium frequency of the residue *α* at position *i* in the presence of drug pressure if only position *i* were allowed to evolve and all remaining residue positions of sequence *S* were held fixed, thus maintaining the background-dependent epistatic constraint on position *i*. Equation 1 gives 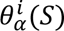 for a specific background *S*, and we also compute the mean adaptive-frequency 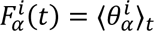 over all sequences in the ensemble at time *t* as the ensemble evolves from the drug naïve to the drug experienced state under the influence of drug pressure. We expect that higher initial mean adaptive-frequency 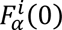 of the DRM in the drug naïve ensemble at *t* = 0 will correspond to a DRM with a faster acquisition rate (i.e. immediately after drug pressure is applied but before additional mutations can accumulate). As a limiting case at some long time after all the residue positions equilibrate under the drug experienced Potts statistical potential, the mean adaptive-frequency must equal the equilibrium frequency of the mutation observed in the Stanford HIVDB of drug-experienced patient sequences, that is: 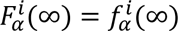 where 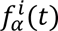 is the frequency of residue *α* at position *i* in the ensemble at time t, i.e. the average over the indicator function 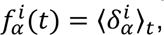 which at *t* = 0 equals the frequency in the drug-naïve dataset and at *t* = ∞ that in the drug-experienced dataset. Note that, 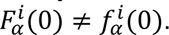

In Fig. 3, we track the time-dependent change in the adaptive-frequency due to epistasis for various DRMs, averaged over an ensemble of trajectories evolving under the drug-experienced Potts statistical potential from the drug-naïve initial state. We simultaneously track the DRM’s frequency over the same ensemble, computed by averaging an indicator function that is 1 when the DRM is present and 0 otherwise. Fitness changes over time as the sequence backgrounds change to accumulate new mutations, and this affects the epistatic constraint on the focal position. As the sequences evolve, the DRM frequency quickly changes to match its adaptive-frequency, which itself changes such that at long times 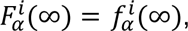 as expected. We focus on a pair of DRMs from each drug target protein. These include: D30N and V32I in HIV-1 protease in response to PIs, where D30N is a “fast” and V32I a “slow” mutation, M184V (fast) and D67N (slow) in RT in response to NRTIs, K103N (fast) and Y181C (slow) in RT in response to NNRTIs, and similarly, N155H (fast) and Q148H (slow) in IN in response to INSTIs. The fast/slow pairs were selected such that their equilibrium frequencies in the initial (drug naïve) and final (drug experienced) states observed in the Stanford HIVDB are approximately the same. We refer the reader to **Table S1-S4** in supporting information for a list of the 52 DRMs that we modeled, together with their drug-naïve and drug-experienced frequencies, and their adaptive-frequency following the application of drugs but before any additional mutations can accumulate.

**Figure 3.**
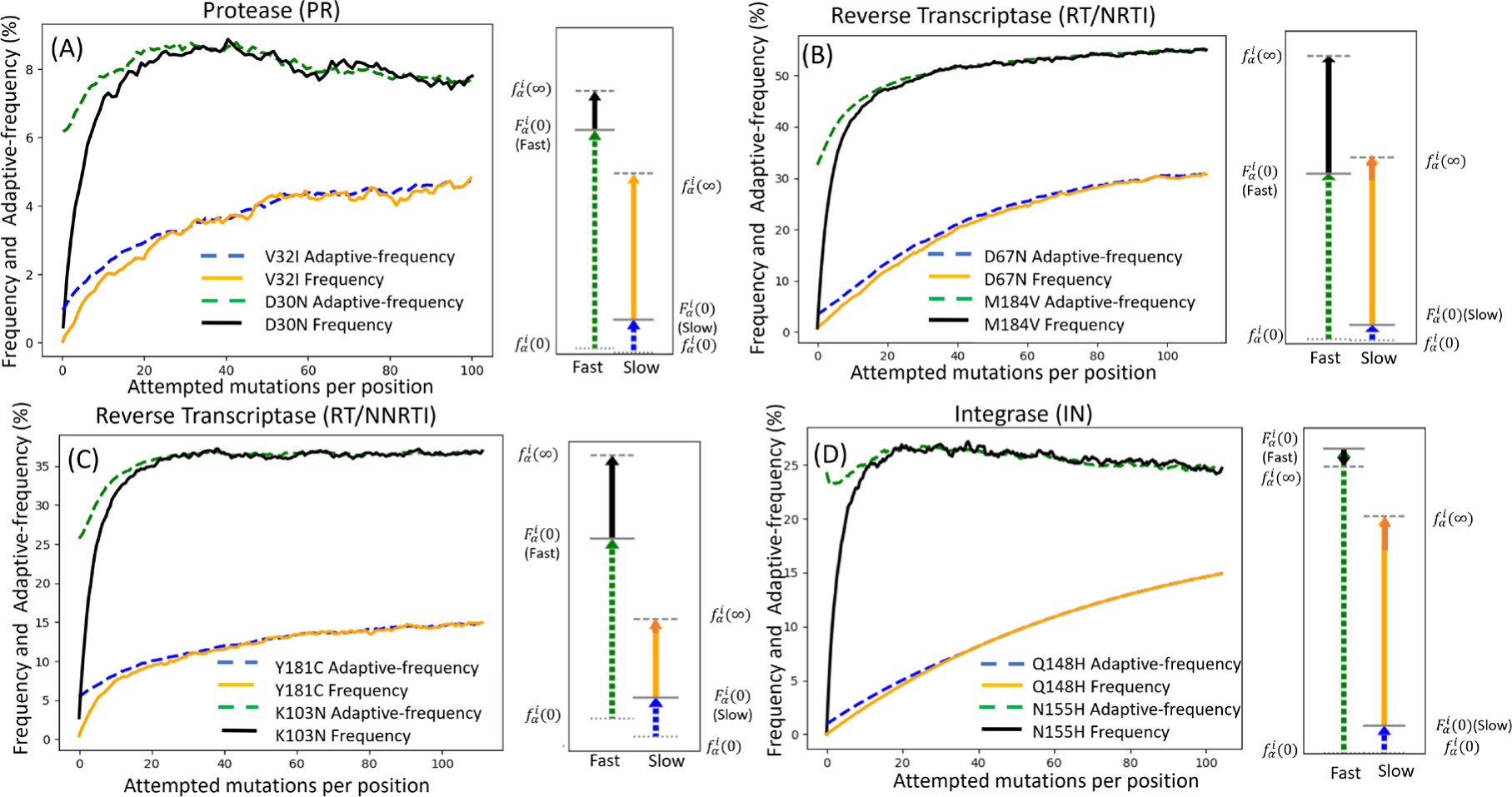
Temporal evolution of the mean adaptive-frequency 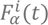 and frequency 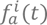 of a mutation as a function of the number of attempted mutations per position for two differently evolving DRMs. The panels refer to (A) D30N (fast) and V32I (slow) DRMs in PR, (B) M184V (fast) and D67N (slow) DRMs in RT in response to NRTIs, (C) K103N (fast) and Y181C (slow) DRMs in RT in response to NNRTIs, and (D) N155H (fast) and Q148H (slow) DRMs in IN in response to INSTIs. The arrows in the right side panel of each plot illustrate schematically the initial adaptive frequency (Initial increase) relative to initial drug-naïve frequency 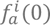 immediately after exposure to drugs at time 0 (dashed green/blue arrow for fast/slow) and the subsequent increase (black/orange solid arrow for fast/slow) from this value to the final value 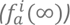 after the background has had time to equilibrate. The initial adaptive-frequency 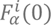 for the faster-evolving DRMs is much closer to the final drug-experienced frequency (dashed green arrows are larger than the dashed blue arrows). The darker orange arrow at the tip of the orange line shows the additional increase in frequency for the slow DRMs after 100 attempted mutations per position. Further discussion about the position-specific contributions of the additional mutations necessary for a focal DRM to occur are discussed in Fig. 4.

The rapidly acquired PR D30N mutation is initially present at less than 1% frequency in the drug naïve ensemble at t=0 but has a significantly higher initial mean epistatic adaptive-frequency of 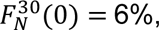 close to its final equilibrium drug-experienced frequency of 8% (**Fig 3A**). Its frequency, i.e., its indicator function average, quickly increases to match its adaptive frequency, and both subsequently equilibrate to their final value of 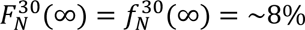 as additional mutations arise in the sequence backgrounds. In contrast, the slowly acquired PR V32I mutation initially has a low mean adaptive-frequency of 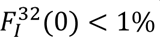 after drug pressure is applied compared to its final equilibrium frequency of ∼5%, and both its frequency and adaptive-frequency slowly rise to this final value. The V32I DRM is slow because additional accessory mutations must co-occur in the sequence background before the DRM’s frequency can become established above the drug naïve level. We refer to this as an *epistatic barrier*, which we find is a central feature of all the DRMs that are acquired slowly. In other words, the slow DRMs are contingent on the appearance of other mutations in the sequence background following the application of drug pressure. The same effects are observed for the M184V (fast) and D67N (slow) for RT/NRTI and the K103N (fast) and Y181C (slow) for RT/NNRTI systems, as illustrated by the examples shown in Fig. 3B**-C**. The IN mutations N155H (fast) and Q148H (slow) are shown as two examples of INSTI-selected DRMs (Fig. 3D) to illustrate the effect of the epistatic barrier on the kinetics of drug resistance for IN.

The results in Fig. 3 show how each DRM’s change in frequency upon drug exposure can be understood as a combination of two effects: (1) the new selective pressures of the drug exposed environment cause an immediate initial increase in DRM adaptive-frequency compared to its frequency in the drug-naïve environment (dashed green/blue arrows in Fig. 3, equal to 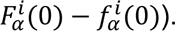 This indicates how, before any other mutations occur in the sequence background, many DRMs would increase in frequency upon exposure to drugs, for instance by providing resistance to the drug directly, even if the stability of the target protein is adversely affected. (2) Subsequently, there is a further increase in adaptive-frequency or fitness mediated through mutations evolving in the sequence background (solid black/orange arrows in Fig. 3, equal to 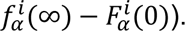 A key distinguishing feature of the fast vs slow DRMs is that the slow DRMs have a small initial increase in adaptive-frequency upon drug exposure (dashed arrows) relative to the overall change (dashed arrow + solid arrow) in that DRM’s frequency at long times due to the combined influence of both the immediate effect of the drug (dashed arrow) as well as fitness contributions from subsequent coupled or accessory mutations (solid arrow). Indeed, we find that the ratio of these two magnitudes, 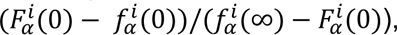 strongly correlates (**Fig. S6, SI**) with the time constant *τ* computed from the kinetic simulations. The fact that this simple ratio computed from the epistatic terms of the Potts model also correlates well with literature-reported DRM acquisition time supports our hypothesis that epistatic coupling is the major determinant of DRM acquisition times.

### Position-specific contributions to the DRM fitness

We next demonstrate how the Potts kinetic model can be analyzed to determine which patterns of mutations lead to an epistatic barrier. Taking advantage of its simple and interpretable form, we define a score to estimate the contribution of each background position to the epistatic barrier for a focal DRM. For a given sequence, we compare the adaptive-frequency for the DRM at position *i* in that sequence to the expected adaptive-frequency if background position *j* were mutated to other residues in proportion to which they appear in the equilibrated (drug-experienced) sequence ensemble. This score is defined as

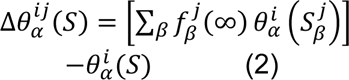

where *i* is the DRM position, *α* the DRM mutation, *j* the background position, and 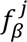 is the frequency of residue *β* at position *j* in the drug-experienced ensemble.

This measures the average change in the sequence-specific adaptive-frequency at position *i* from its initial value caused by letting only position *j* evolve to the drug-experienced ensemble frequencies, keeping all other residues in the sequence fixed. In this way, only positions *j*, which are both epistatically coupled to position *i* and have mutations *β* that arise during evolution to the drug experienced ensemble, will have significant Δθ(*S*) values. Further, we average 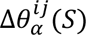 over the drug-naïve sequence ensemble to give a "change in fitness" score 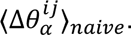 The sum of this score over all positions 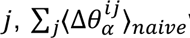 will approximate the net change in adaptive-frequency of the DRMs at position *i* due to its direct coupling to all positions *j* as the ensemble evolves under drug pressure to its drug-experienced values (solid arrows).

The coupled mutations identified with equation (2) are shown in Fig. 4 for pairs of fast and slow DRMs from each protein target. The additional coupled mutations are largely consistent with the literature, in that numerous studies identify them as being associated with each focal DRM. This observation shows that the Potts predicted epistatic barrier can be rationally decomposed and additionally implies that such analyses can be used to quantitatively identify and study novel couplings that were previously missed. **Section S1** of the supporting information contains a detailed discussion about the literature survey pertaining to every pair of fast and slow mutations featured in Fig. 4, along with an additional discussion on the RT/NRTI Q151M complex (**Fig. S7**, **SI**). This decomposition is not only broadly consistent with the literature but also quantifies the relative strength of the coupled interactions for each DRM and specifically identifies “directly” coupled positions as opposed to mutations indirectly coupled through an epistatic network.

**Figure 4:**
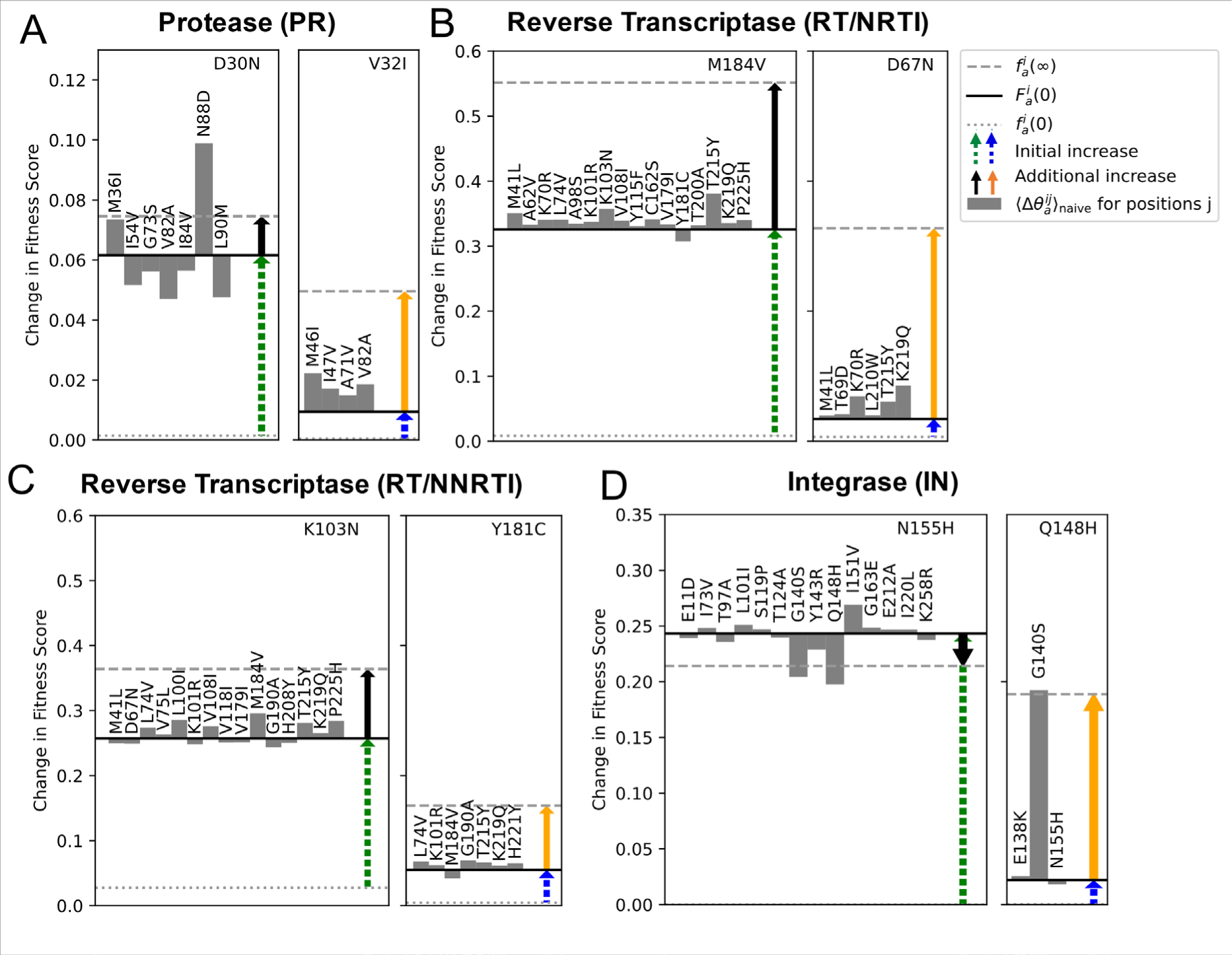
Comparison of the additional mutations necessary for a focal DRM to occur when comparing DRMs with “fast” and “slow” acquisition times. The DRMs shown are (A) PI DRMs in PR, for D30N (fast) and V32I (slow), (B) NRTI DRMs in RT for M184V (fast) and D67N (slow), (C) NNRTI DRMs in RT for K103N (fast) and Y181C (slow), and (D) INSTI DRMs in IN for N155H (fast) and Q148H (slow). In each plot, the focal DRM is listed in the upper right. The dotted grey line reflects the DRM’s frequency in the drug-naïve ensemble, the solid black line reflects the DRM’s average fitness under the drug-experienced Hamiltonian in the drug-naïve ensemble, and the dashed grey line reflects the DRM’s frequency in the drug-experienced ensemble. The dashed green/blue arrow then represents the initial increase in the DRM’s frequency upon exposure to drugs starting from the drug-naïve state if the background were held fixed, identical to the dashed green/blue arrows in figure 3, and the solid black/orange arrow represents the additional increase in DRM frequency once the sequence backgrounds are allowed to vary and accumulate coupled mutations, identical to the solid black/orange arrow in figure 3. The grey bars represent the fitness measured using equation 2, averaged over the drug-naïve ensemble, giving the first-order contribution to the increase in the fitness of the DRM at i due to the evolution of j to the drug-experienced state. The sum of the grey bar magnitudes approximates the length of the solid black/orange arrow. For the fast DRMs, the effect of the initial increase in fitness (dashed green arrow) is larger than the additional increase in fitness due to coupled mutations (dashed blue arrow).

### Structural Underpinnings for the rate of emergence of DRMs

Our study highlights the central role epistasis plays in kinetics leading to major DRMs in HIV, which can span a large time range. However, these predictions do not explain the mechanistic origins for different rates of emergence of DRMs. Here we explore a structural rationale for the differential fitness of resistance mutations in a drug naïve background under drug selection pressure. We include examples of fast and slow mutations from each of the four drug classes. The analyses suggest a general principle whereby faster mutations induce changes that are less disruptive and can be more readily compensated, whereas slower mutations typically induce more disruptive changes and/or lead to drug excision through indirect mechanisms.

Fig. 5A shows the changes in the PR active site evolving in response to PIs. The fast-arising mutation D30N leads to the loss of a crucial hydrogen bond formed between the inhibitor and PR, which immediately reduces binding affinity and engenders drug resistance. Although the removal of negative charge also results in altered electrostatic interaction with the native substrate, and reduced proteolytic activity, this is easily overcome by the compensatory charge swap mutation N88D, which also arises quickly (48). D30N, in conjunction with N88D, rapidly emerge and mutually entrench each other. In contrast to the fast-arising D30N (and N88D), the slow-arising mutation V32I works through an indirect mechanism. V32 resides at the periphery of the PR active site and makes an important network of hydrophobic interactions with residues I47 and V82 that frequently arise in concert with V32I, all of which are required for PR catalytic activity (49). The mutation V32I causes an extensive pattern of rearrangements that ultimately results in repositioning of the inhibitor. PR variants containing V32I mutations frequently display comparatively larger dynamic fluctuations, which propagate throughout the enzyme, in comparison to other PR variants. For example, as many as 12 hydrogen bonds can change significantly in comparison to the WT enzyme (50). Thus, the fast-arising mutation D30N directly affects ligand binding, whereas V32I works through an indirect mechanism.

**Figure 5:**
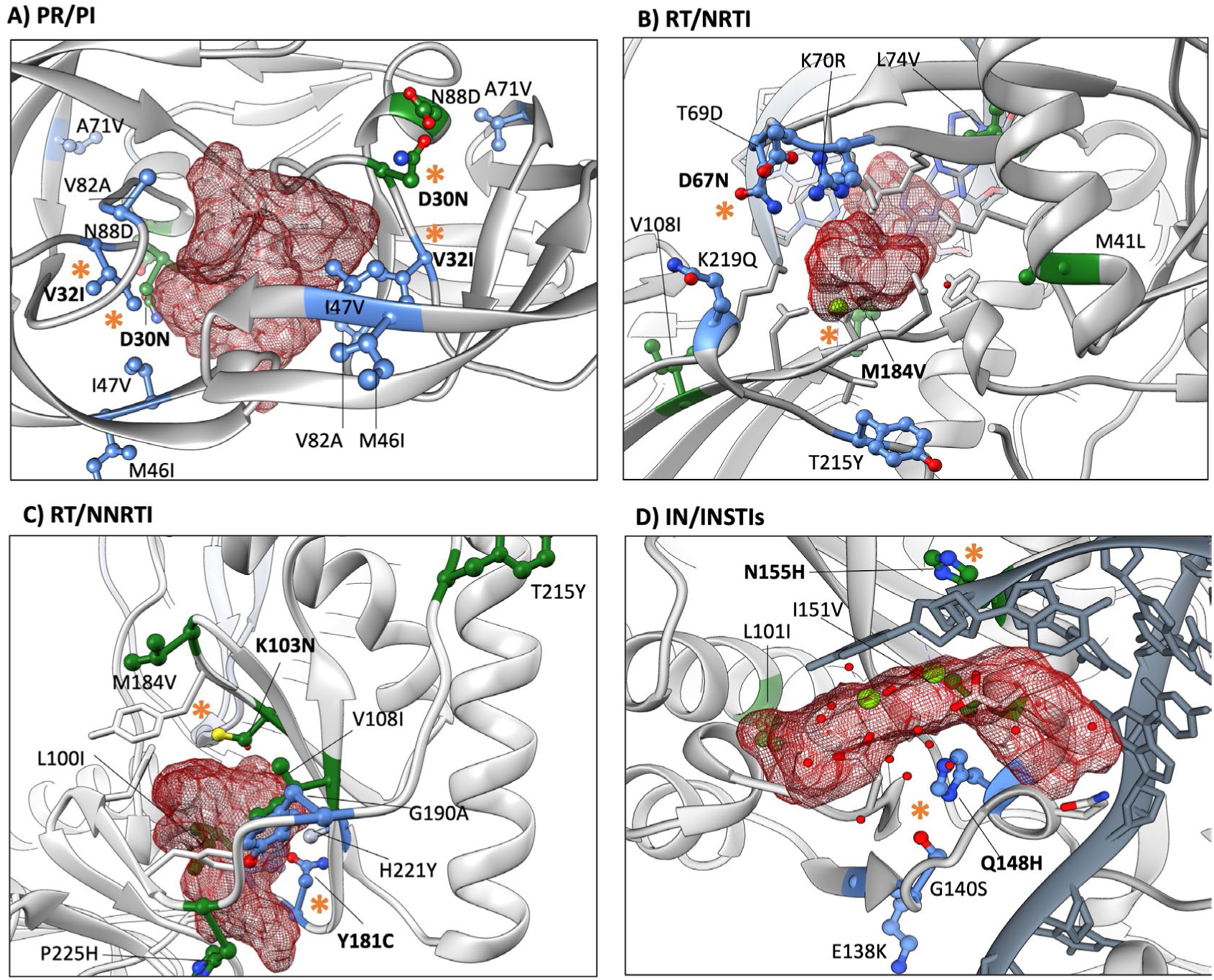
The structural underpinnings for the rate of emergence of DRMs are shown for all four classes of drugs. Fast (green) and slow (blue) mutations are shown for (A) HIV-1 PR against PIs (PDB: 7DPQ, 4Q1X), (B) RT against NRTIs in (PDB: 6UJY), (C) RT against NNRTIs (PDB: 3BGR), and (D) IN against INSTIs (PDB: 8FNP). The extent of the known molecular envelope for different bound drugs is shown as a red mesh (More details in the computational method section). The slow and fast mutations being followed are indicated in bold and orange asterisks (*). The other mutations involved in pathways with the slow or fast mutations are shown in the same color code (green and blue). Where mutant structures are not available, mutations are introduced through the UCSF Chimera (60) Structure Editing module and the highest probable rotameric configurations of the mutation side chains, based on previously determined atomic structures, are shown.

Fig. 5B shows the changes in the RT active site evolving in response to NRTIs. The fast-arising M184V DRM in RT results in fewer disruptive changes to the enzyme and acts directly to displace bound ligand. All approved NRTIs lack a 3-OH and, when incorporated into the nascent DNA primer strand by RT, act as chain terminators. The M184V mutation replaces a flexible side chain near the polymerase active site with a branched amino acid that selectively discriminates against NRTIs, while still allowing for the incorporation of dNTPs with normal deoxyribose rings. Thus, M184V directly displaces the NRTI, but has minimal effect on normal enzyme activity. The slow-arising D67N mutation, however, resides in a different location – at the tip of the flexible 3-4 loop with its side chain facing the ATP – and typically arises in combination with other mutations. D67N retains a similar size but eliminates the negatively charged environment imparted by the original Asp67. To compensate for this change in the electronic environment, RT must acquire additional subtle and interconnected background mutations that would remain conducive to ATP binding while allowing the mutant enzyme to excise a broad array of NRTIs (51). Thus, D67N arises more slowly, due to the requirement of developing compensatory background mutations that increases the fitness substantially above the drug naïve value with the D67N mutation.

Fig. 5C shows the changes in the RT active site evolving in response to NNRTIs. The fast-arising K103N DRM leads to a novel hydrogen bond between N103 and Y188, which is otherwise absent in WT RT. The protein interaction network surrounding the newly formed hydrogen bond stabilizes the closed-pocket conformation of the enzyme, thus impeding NNRTI access to the binding pocket (52, 53). Notably, K103N induces minor changes to the pocket residues compared to WT RT. However, resistance due to the slow-arising Y181C is more disruptive to the binding pocket. Y181C abrogates the π–π stacking interactions between two aromatic rings of residues on RT (Y181, Y188) and an aromatic ring of bound NNRTIs (52, 54, 55). Changes associated with the binding pocket in RT are more extensive in response to Y181C mutations, and thus the enzyme benefits from the addition of specific additional background mutations. Again, the slow development of Y181C can be explained by the requirement to develop additional compensatory changes.

Fig. 5D shows the changes in the IN active site evolving in response to INSTIs. The fast-arising mutation N155H leads to a salt-bridge interaction with the vDNA phosphate, which was hypothesized to affect the kinetics of INSTI binding (56). Enzyme activity is minimally affected by the N155H mutation. In contrast, the slow-arising mutation Q148H is well-known for its detrimental effect on enzyme activity. The mechanism of resistance for Q148H can be explained by the introduction of an electropositive moiety underneath the two Mg^2+^ metal ions, weakening metal chelation and leading to INSTI displacement (57, 58). Importantly, Q148H also significantly compromises enzyme activity, because the Mg^2+^ ions are also directly involved in catalysis. Since Q148H leads to a more extensive modulation to the structure by itself, this DRM also leads to a more substantial fitness cost in the drug naïve viral population than N155H. To account for the greater drop-in fitness associated with Q148H, the key compensatory G140S mutation must evolve to restore replicative capacity, while other mutations frequently accumulate with this G140S/Q148H pair. Despite the very different timescales for DRM emergence, both N155H and Q148H are the two most frequently encountered IN mutations in the Stanford drug resistance database (59), indicating that the rate of emergence of the DRM is not necessarily correlated to its final frequency in the population; the final frequencies depend heavily on the background.

To generalize the analyses from structural biology, there are two scenarios that discriminate fast vs. slow mutations in the context of drug binding, which may affect either [1] ligand binding or [2] enzymatic activity. In the first scenario, assuming enzymatic activity remains constant, fast mutations will affect the ligand binding directly, whereas slow mutations may work through indirect mechanisms and must accumulate in conjunction with other changes that eventually displace the ligand. In the second scenario, assuming that their effect on ligand binding remains constant, fast mutations will generally have a smaller effect on enzyme fitness, whereas slow mutations lead to more profound detrimental changes that affect the natural function of the enzyme and must therefore be compensated by additional background changes. We note that these scenarios are not mutually exclusive, and most cases are likely to be explained by a combination of these effects.

## Conclusion

The evolution of HIV under drug pressure and internal epistatic constraints induces correlated mutations that change the frequencies at which DRMs appear in the population over time. Literature surveys show that the timeline of emergence of DRMs from the drug naïve patient population varies from a few weeks to a year or more when these patients receive ART. In the present study, we modelled the kinetics of the emergence of DRMs using Kinetic Monte-Carlo simulations on a fitness-landscape described by an epistatic Potts statistical potential parameterized on drug-experienced sequence data. We propagate an initial sequence ensemble that matches the patterns observed in the drug-naïve population as it evolves to a final ensemble that matches the patterns observed in the drug-experienced ensemble.

We selected 52 DRMs from three different protein targets (PR, RT [NRTI and NNRTI], IN) to study their kinetics by calculating the acquisition times (τ) and compared them with the timeline of emergence reported in the literature. The times to acquire drug-resistance mutations predicted by the KMC model are highly correlated with the acquisition times reported in the literature (ρ = ∼0.85, p<<0.001). Qualitatively, for DRMs that are reported in the literature as acquired rapidly (emergence time range between 0.5—3 months), the predicted KMC time constants are *τ* < 10; while for DRMs reported in the literature as acquired slowly (∼8—20 months), the KMC time constants are *τ* > 24. These results provide strong evidence in support of the role of epistasis – the couplings between DRMs – as the determining factor, which distinguishes DRMs that are acquired rapidly from those that are acquired slowly.

We introduced a metric, the sequence dependent adaptive-frequency 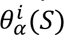 as a proxy for fitness, which is a measure of the likelihood of a DRM in a fixed background *S* under the drug-experienced Potts statistical energy model, and the corresponding ensemble averaged adaptive-frequency 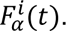 The most important feature that distinguishes DRMs that are acquired slowly from those that are rapidly acquired is the initial fitness of the DRM in the sequence when drug pressure is applied before additional mutations have time to accumulate. For DRMs acquired rapidly, the frequency begins to increase within the drug-naïve ensemble as soon as drug pressure is applied, without the need to first accumulate additional mutations. In contrast, for DRMs acquired slowly, the frequency increases in tandem with the accumulation of additional mutations. We interpret this as the existence of an “epistatic barrier” for the slow DRMs, consistent with results suggesting epistasis reduces evolutionary rates (35, 36). In contrast, two other factors that have been suggested to have a large effect on the rates at which DRMs are acquired, including within host effects associated with the genetic code (transitions (Ti) vs. transversions (Tv), and the number of nucleotide changes (Δnuc) required per codon change) and the overall fitness of the DRM as estimated by its prevalence in the drug-experienced population, are not well correlated with the times to acquire drug resistance reported in the literature.

This work provides a framework for the development and application of computational methods to forecast the time course and the pathways over which drug resistance to antivirals develops in patients. While the KMC simulations for each of the three HIV target enzymes (PR, RT, and IN) consist of tens of thousands of trajectories that propagate the initial drug naïve ensemble to the final drug-experienced ensemble, both of which are observed in the Stanford HIVDB, we expect that there are a much smaller number of pathways that can be discerned by clustering the trajectories and identifying structural constraints that can be used to distinguish the clusters and annotate them. These ideas are the subject of ongoing research.

### Computational methods

In this section, we present the Potts Hamiltonian model and the motivation behind the model. We also describe the kinetic Monte Carlo methods which are used to study the temporal evolution of HIV under drug selection pressure (34, 60). Here, we measure time in these kinetic simulations in the units of the number of mutations attempted per position, i.e. the number of attempted mutations throughout the sequence divided by the sequence length. The length of PR, RT, and IN are 99, 188 and 263 respectively (33); and with this normalization scheme of the KMC algorithm ensures that, a time unit has the same meaning across the three HIV enzymes, PR, RT, and IN, which have differential lengths. The details of the data processing, mutation classification, alphabet reduction and length of amino acids are discussed in our previous paper (34).

The UCSF Chimera (61) Structure Editing module is used to prepare Fig. 5. The extent of the known molecular envelope for different bound drugs is shown as a red mesh; The molecular envelope is created using UCSF Chimera (58) and using the PDBs: PIs: DRV (PDB: 4Q1X) SQV (PDB: 3S56), NFV (PDB: 3EKX), DRV (PDB: 3D20), LPV (PDB: 2QHC), TPV (PDB: 4NJU); NRTIs: FTC (PDB: 6UIR), AZT (PDB: 3KLG); NNRTIs: RPV (PDB: 3BGR), DOR (PDB: 7Z2G), EFV (PDB: 1IKW), ETR (PDB: 3MEC), NVP (PDB: 4PWD); INSTIs: DTG (8FNZ), BIC (6PUW).

### Potts Hamiltonian model

We use the Potts model which is a probabilistic model designed to describe the probabilities of observing specific states of a system that is constructed to be as unbiased as possible except to agree with the average first- and second-order observable (marginals) from the sequence data (62-68). The Potts model has a long history of use in statistical physics and analysis of protein sequence. In a set of protein sequences, the single and pair amino acid frequencies are average quantities that can be estimated from the finite samples using the data. The details of the models are described in our previous work (34, 37).

### Kinetic Monte Carlo (KMC) simulations

The kinetic Monte Carlo simulation is a Monte Carlo method which is intended to simulate the time evolution processes with known transition rates between states.

The Metropolis algorithm (69) is used to evaluate the metropolis acceptance probability of a mutation such as W (Wild-Type)→M (Mutant) at a randomly chosen position i in a given sequence background at every simulation step given by 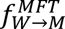

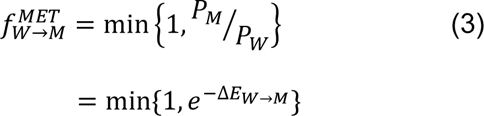

where ΔE_W →M_ = E_M_ – E_W_ is the change in Potts energy in going from residue W to M in the given background.

At the beginning of the simulation process a set of seed sequences (Drug naïve sequences) are taken and a random position *i* and random mutant residue α (Reduced alphabet of 4 letters are used) are chosen the amino acid character at the chosen position i is either preserved or mutated based at the chosen position i is either preserved or mutated based on based on the Metropolis acceptance rate 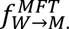 For example, a mutation V32I from V (valine) → I(isoleucine) at position 32 in HIV-1 protease (99 residues long) has a probability 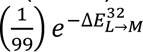 associated with the mutation at each KMC step. The algorithm used here allowed self-mutations during the simulation.

Our kinetic Potts model is a coarse-grained representation of the within-host evolutionary process. Evolution within a host is driven by various selective forces due to host immune response or to maintain viral viability, non-selective forces such as genetic drift, and is affected by other well-known aspects of HIV infection such as retroviral dormancy, compartmentalization, and high recombination rates. In the chronic phase of HIV infection, a large number of viral variants may be present at any point in time, however this population is typically measured to have high sequence identity of close to 99% between pairs of viral genomes in a single host. This is much larger than the typical sequence identity of consensus sequences from different hosts of 90% for our datasets and justifies summarizing a host’s viral population by a single “consensus” sequence. Additionally, the host consensus sequence is observed to accumulate substitutions in a relatively clock like manner, suggestive of sequential selective sweeps. Therefore, instead of modelling the detailed “microscopic” evolutionary forces we use a coarse-graining which only tracks the consensus sequence of a host viral population over time as it accumulates substitutions due to these underlying forces.

In this way, one interpretation of our coarse-grained kinetics is that it models a series of point-mutation events in a viral population which occur according to a Poissonian mutational process, and these mutations are either fixed or lost from the population according to a fitness landscape inferred based on between-host sequence data. We coarse-grain a number of aspects of evolutionary dynamics, for instance we model amino-acid sequences instead of nucleotide sequences, assuming all amino acids can mutate to all others as is commonly done in phylogenetic analyses, for instance in the WAG and JTT models. While this coarse-grained model is necessarily a simplification of HIV viral dynamics, there are key properties of its construction which support conclusions drawn from it. First, an inferred “Potts” model prevalence landscape will implicitly capture many of the averaged effects of various microscopic evolutionary forces because it is fitted to HIV sequence datasets which arose under the microscopic dynamics. For instance, it will capture mutational biases as these causes an increase in the inferred prevalence of the biased amino acids. Second, this model is numerically consistent with the observed between-host sequence variation data: If we use this kinetic model to simulate parallel trajectories (representing evolution in different hosts) and collect the final sequences, then by construction the mutational statistics of the generated sequences (frequencies of amino acids and k-mes of amino acids) match those of the between-host sequence datasets used to train the fitness model. We use a particular inference technique which we have confirmed gives a generative model which very closely reproduces the natural patterns of HIV sequence variation for high order k-mes in generated sequences.

We assume an underlying Poisson mutational process, such that mutation arises at a rate µ. We implemented this by assigning each step in the Metropolis Algorithm a time drawn from an exponential distribution which is the waiting time for a Poisson process.

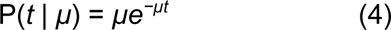

With this overall KMC scheme our simulations match empirical data in two ways. First, the value of µ can be calibrated so that the simulated evolutionary trajectories accumulated substitutions at the same rate as observed experimentally. Second, when using this scheme to run many parallel trajectories until equilibrium, the bivariate residues of the resulting MSA match those observed in the HIV sequence database used to train the Potts model.

### Potential coarse graining approximations

Here we discuss possible coarse-graining errors we have investigated. One aspect which our model has coarse-grained is drug-specific selection pressure and host-specific immune pressure. The ranges for literature acquisition times listed in table 1 reflect, in part, the fact that some mutations arise in response to multiple drugs, but with some difference in acquisition time for each drug, this effect cannot be captured by our fitness model which reflects an averaged selection pressure of all drugs. But the overall correspondence of our model and literature suggests the major determinant of acquisition time is the epistatic interaction of the primary DRM with accessory mutations, which is captured by our fitness model and is independent of specific drug. It may be possible to explicitly model such drug-specific modulation of selection strengths through extensions of our fitness model, which we intend to investigate in the future.

Our model also coarse-grains the mutational process, and for instance does not explicitly distinguish between transition (Ti) and transversion (Tv) mutations, which occur at different rates, and does not explicitly distinguish between mutations in the amino-acid sequence which correspond to single-nucleotide and double-nucleotide mutations at the nucleotide level. Because our fitness landscape is inferred from HIV sequences which evolved in vivo under the influence of these mutational biases, the model implicitly captures their effect to some degree. To further investigate whether these mutational biases significantly affect DRM acquisition time, we investigated their effect for the mutants listed in Table 1. The alterations in the genetic code corresponding to each DRMs are listed in the second column of **Table 1** (**Table S1-S4** in Supporting Information) to assess the impact of genetic codes on the timeline of drug resistance evolution. The majority of DRMs, regardless of the protein type (PR, RT, or IN), are linked to single nucleotide changes (approximately 90%). Consequently, for these mutants, the number of genetic code alterations does not directly affect the time required to develop drug resistance in HIV. The DNA substitution mutations can be categorized as transition (Ti) or transversion (Tv). Transitions involve the exchange of two-ring purines (A → G) or one-ring pyrimidines (C → T), representing bases with similar sizes. On the other hand, transversions involve the interchange of purine for pyrimidine bases, which entails the exchange between one-ring and two-ring structures. We have included the nature of nucleotide changes (Δnuc) for each DRM of all proteins (PR, RT, and IN) in the third column of **Table 1** (**Table S1-S4** in Supporting Information). Our model does not provide information about the DNA level of the sequences; therefore, silent substitutions due to wobble base pair effects are not considered when determining the nature of nucleotide changes. In summary, we conclude that mutational biases have a negligible or relatively minor influence on acquisition time, whereas the epistatic interactions captured by our fitness model have a much more significant effect.

## Data Availability

All study data are included in the article and/or supporting information.

## Supporting Information

This article contains supporting information online. The supporting information includes the following: (1) The distribution of the number of inhibitor or drug-associated mutations in HIV-1 in PR and IN (Fig. S1 and S2). (2) Evolution of drug resistance from drug-naïve state to drug-experienced state in RT (NRTI), PR and IN (Fig. S3). (3) Acquisition times of emergence of major DRMs in IN, RT (NRTI) and PR (Table S1-S3), (4) The Spearman rank correlation between the literature timeline and the Potts+KMC simulation predicted time constants (τ) (Fig. S4), (5) Fitness across sequence backgrounds for primary drug resistance mutations in PR, RT (NRTI+NNRTI) and IN (Table S4-S7), (6) The effect of epistasis on the favorability of a primary resistance mutation in protease (PR) and reverse transcriptase (RT) (Fig. S5), (7) Spearman rank correlation between the new drug-exposed environment to the initial increase in adaptive frequency and the Potts+KMC simulation predicted time constants (τ) in different protein targets (Fig. S6), (8) Comparison of the additional mutations necessary for Q151M (RT/NRTI) to occur (Fig. S7). (9) Summary of literature evidence of the impact of identified accessory mutations on the kinetics of "Fast" and "Slow" drug resistance mutations (Section S1).

## Acknowledgement

This work has been supported by the National Institutes of Health through grants awarded to Ronald M. Levy (U54-AI150472, R01 AI178849, S10OD020095). The National Science Foundation also provided funding through a grant awarded to RML and AH (1934848). EA was supported by NIH grants U54 AI150472 and R01 AI027690. DL was supported by NIH grants U54 AI150472, U01 AI136680, R01 AI146017, the Margaret T. Morris and the Hearst Foundations. The funders had no role in study design, data collection and analysis, decision to publish, or preparation of the manuscript.

## Supporting Information

**Figure S1:**
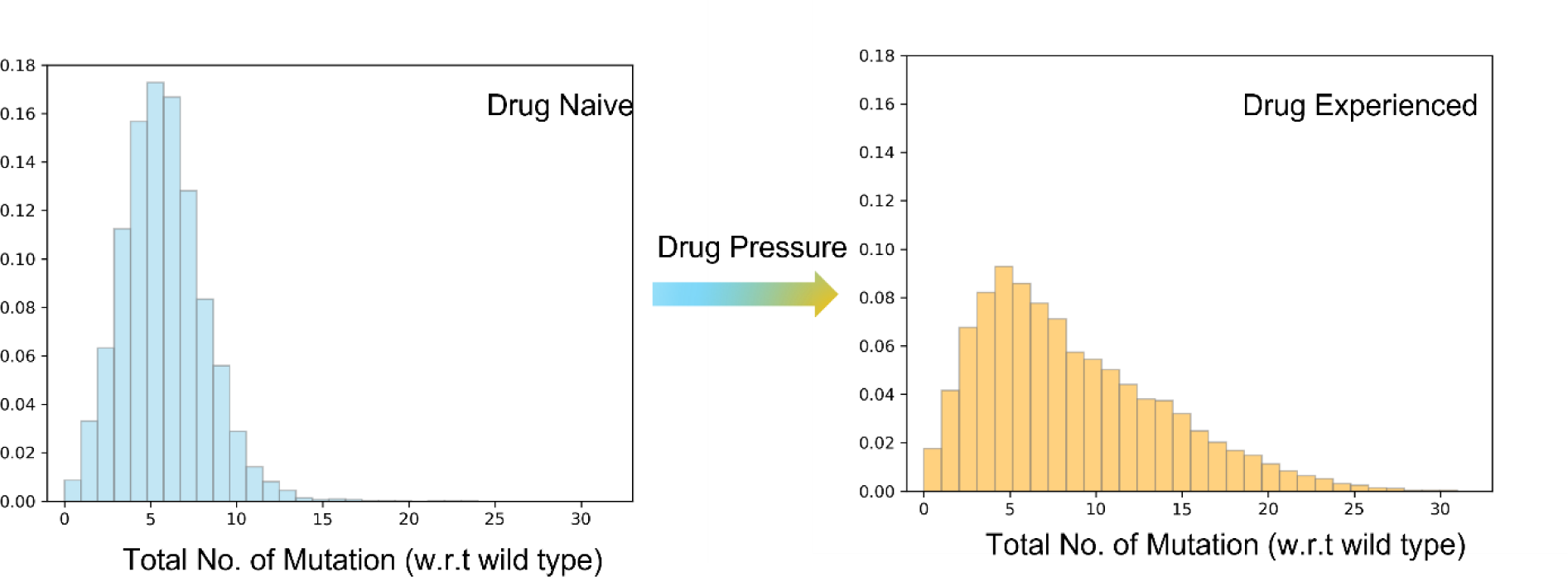
The distribution of the total number of mutations in HIV-1 protease (PR) in (A) the drug-naive multiple sequence alignment (MSA) available from the Stanford HIV database, and (B) the drug-experienced MSA (1).

**Figure S2:**
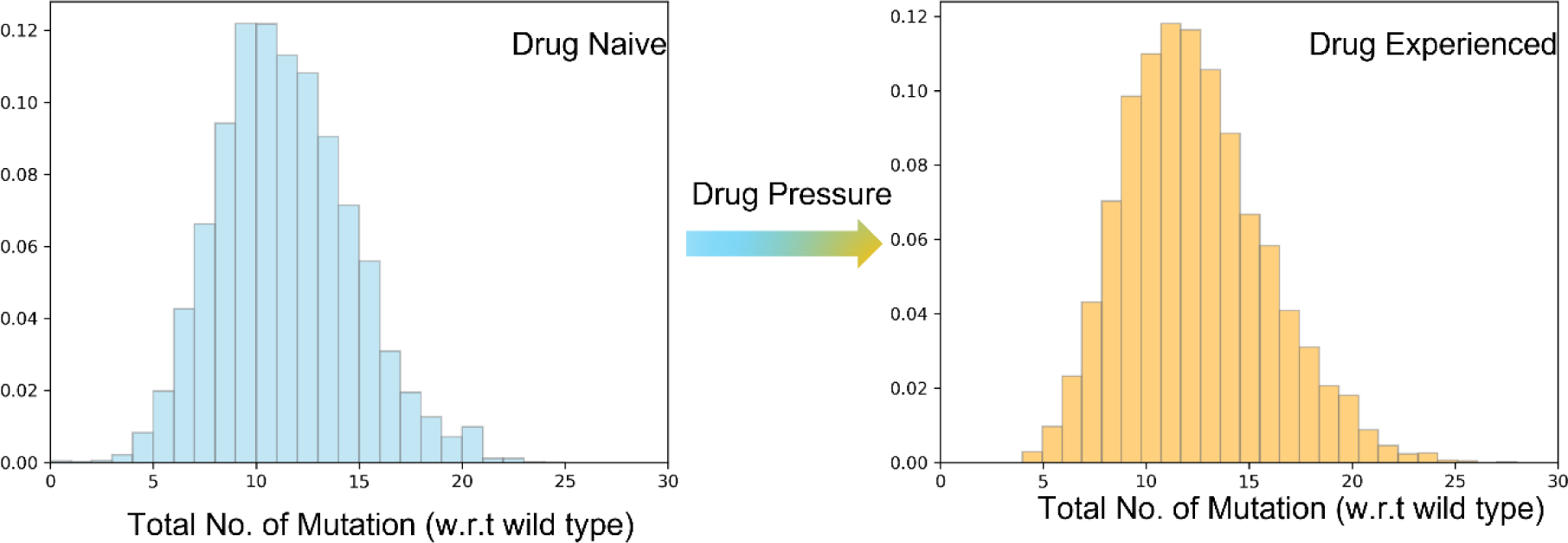
The distribution of the total number of mutations in HIV-1 integrase (IN) in (A) the drug-naive multiple sequence alignment (MSA) available from the Stanford HIV database, and (B) the drug-experienced MSA (1).

**Figure S3:**
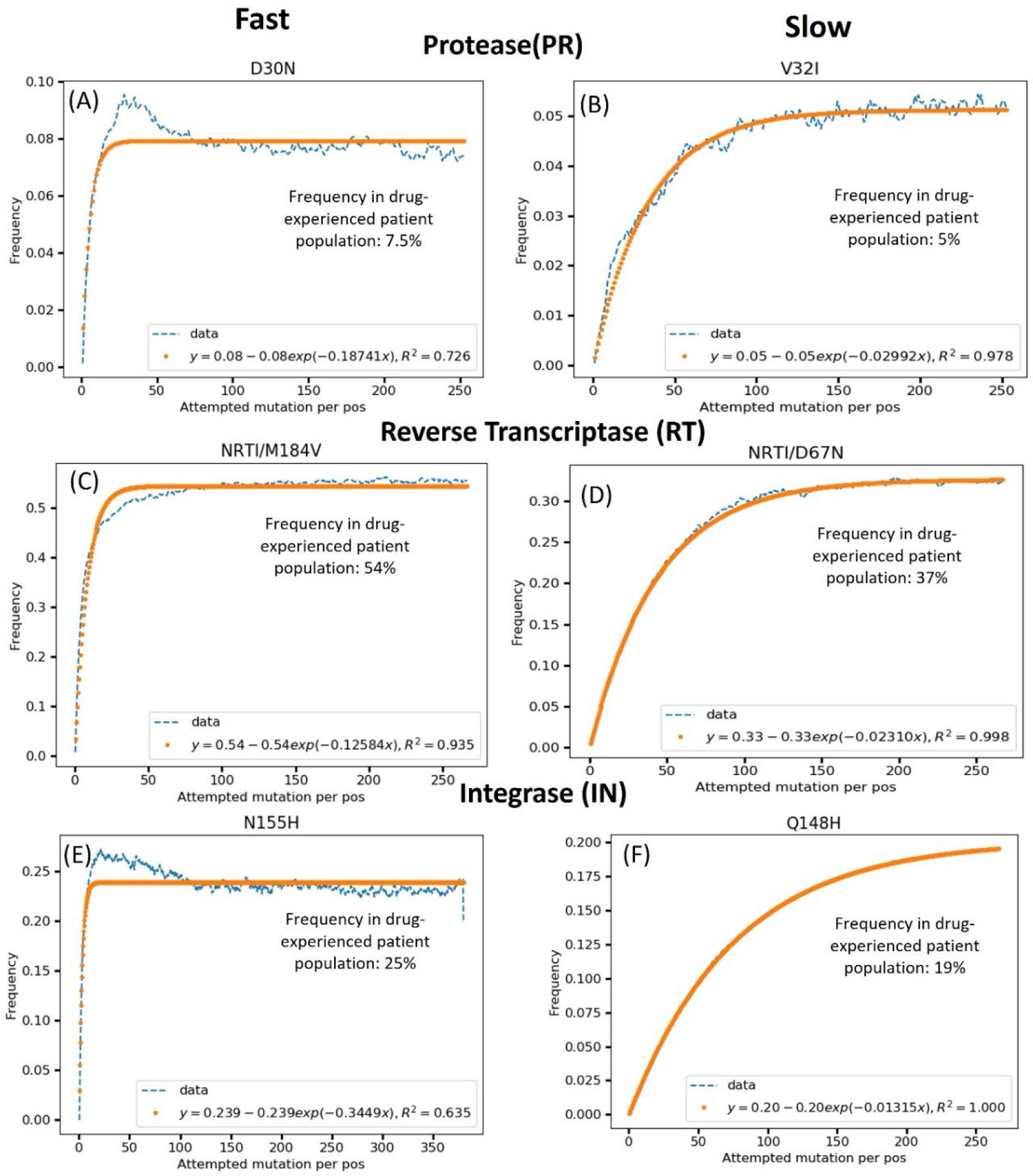
Evolution of drug resistance from the drug-naïve state to drug-experienced state in protease (PR), reverse transcriptase (RT/NRTI), and integrase (IN). The change in frequencies of primary drug resistance mutations (A) D30N, (B) V32I in in protease (PR) (upper panel), (C) M184V, (D) D67N in reverse transcriptase (RT/NRTI) (middle panel) and (E) N155H, (F) Q148H integrase (IN) (lower panel). The simulation was started with drug-naïve MSA from the Stanford HIV Database (1) and evolved towards drug-experienced state. Evolution time is represented as a function of the number of mutations attempted mutations per position (the length of PR is 99, RT is 188, and IN is 263).

**Table S1.** Acquisition times of emergence of major protease (PR) resistance mutations using KMC simulation and literature survey, which can occur as a transition (Ti), transversion (Tv) or both, and involve different numbers of nucleotide changes (Δnuc). The table is ordered by the measured by the estimated time constant **τ**.

| Mutations | $\Delta\text{nuc}$ | Nature of Mutation | Frequency (%) | | | Time Constant ($\tau$) | Timeline in Literature (Months) | Classification of Timeline (Literature) | Reference |
| --- | --- | --- | --- | --- | --- | --- | --- | --- | --- |
| | | | Drug-naive | Drug-experienced | Adaptive frequency $\langle F_a^i(t=0) \rangle$ | | | | |
| D30N | 1 | Ti | 0.14 | 8 | 6.21 | 5.33 | ~ 1-3 | Fast | (2, 3) |
| N88D | 1 | Ti | 0.12 | 5 | 2.9 | 9.02 | ~ 1-3 | Fast | (3, 4) |
| L90M | 1 | Tv | 0.78 | 31 | 10.4 | 25.51 | ~ 6-12 | Slow | (3, 5, 6) |
| M46I | 1 | Ti | 0.42 | 21 | 5.10 | 25.86 | ~ 5-9 | Slow | (5, 7) |
| I50V | 1 | Ti | 0.026 | 1.0 | 0.05 | 27.03 | ~ 10-11 | Slow | (8) |
| V82A | 1 | Ti | 0.26 | 24 | 4.20 | 28.80 | ~ 8-11 | Slow | (9, 10) |
| G48V | 2 | Tv | 0.01 | 5 | 0.7 | 32.23 | ~ 6.5-11.5 | Slow | (10, 11) |
| I84V | 1 | Ti | 0.17 | 13 | 0.3 | 32.50 | ~ 10-12 | Slow | (5, 12) |
| V32I | 1 | Ti | 0.04 | 5 | 0.1 | 33.42 | ~ 6-12 | Slow | (10, 12-14) |
| L76V | 1 | Tv | 0.02 | 3 | 0.5 | 32.90 | ~ 9 | Slow | (15) |
| I54V | 1 | Ti | 0.22 | 23 | 2.1 | 33.81 | ~ 9-10.5 | Slow | (5, 10) |
| I47V | 1 | Ti | 0.7 | 4 | 0.7 | 41.44 | ~ 9-13 | Slow | (16, 17) |
| I54L/M | 1 | Tv | 0.066 | 4 | 0.6 | 43.85 | ~ 10-11 | Slow | (17) |

**Table S2:**
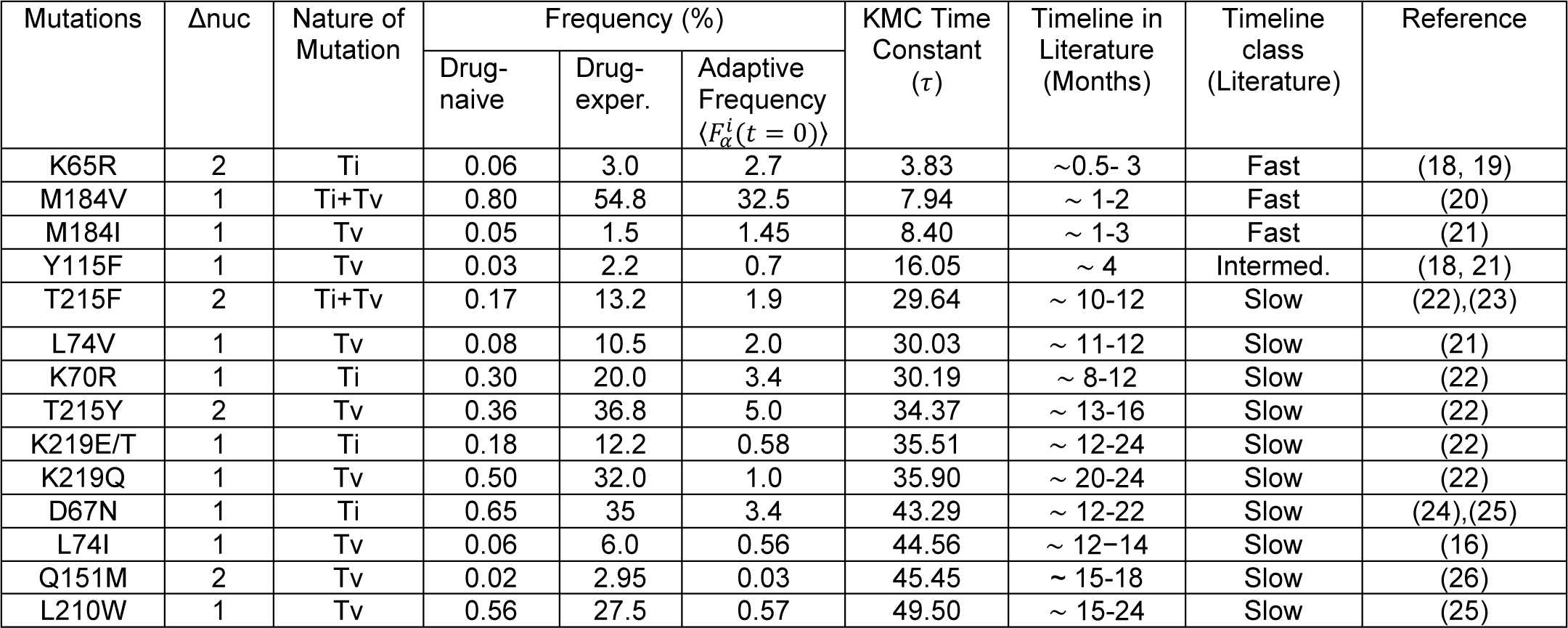
Acquisition times for nucleoside-analog RT inhibitor (NRTI) selected primary resistance mutations using KMC simulations and the literature survey, which can occur as a transition (Ti), transversion (Tv) or both, or can involve different numbers of nucleotide changes (Δnuc). The table is ordered by the measured by the estimated time constant **τ**.

| Mutations | $\Delta\text{nuc}$ | Nature of Mutation | Frequency (%) | | | KMC Time Constant ($\tau$) | Timeline in Literature (Months) | Timeline class (Literature) | Reference |
| --- | --- | --- | --- | --- | --- | --- | --- | --- | --- |
| | | | Drug-naive | Drug-exper. | Adaptive Frequency $\langle F_a^i(t=0) \rangle$ | | | | |
| K65R | 2 | Ti | 0.06 | 3.0 | 2.7 | 3.83 | ~0.5- 3 | Fast | (18, 19) |
| M184V | 1 | Ti+Tv | 0.80 | 54.8 | 32.5 | 7.94 | ~ 1-2 | Fast | (20) |
| M184I | 1 | Tv | 0.05 | 1.5 | 1.45 | 8.40 | ~ 1-3 | Fast | (21) |
| Y115F | 1 | Tv | 0.03 | 2.2 | 0.7 | 16.05 | ~ 4 | Intermed. | (18, 21) |
| T215F | 2 | Ti+Tv | 0.17 | 13.2 | 1.9 | 29.64 | ~ 10-12 | Slow | (22),(23) |
| L74V | 1 | Tv | 0.08 | 10.5 | 2.0 | 30.03 | ~ 11-12 | Slow | (21) |
| K70R | 1 | Ti | 0.30 | 20.0 | 3.4 | 30.19 | ~ 8-12 | Slow | (22) |
| T215Y | 2 | Tv | 0.36 | 36.8 | 5.0 | 34.37 | ~ 13-16 | Slow | (22) |
| K219E/T | 1 | Ti | 0.18 | 12.2 | 0.58 | 35.51 | ~ 12-24 | Slow | (22) |
| K219Q | 1 | Tv | 0.50 | 32.0 | 1.0 | 35.90 | ~ 20-24 | Slow | (22) |
| D67N | 1 | Ti | 0.65 | 35 | 3.4 | 43.29 | ~ 12-22 | Slow | (24),(25) |
| L74I | 1 | Tv | 0.06 | 6.0 | 0.56 | 44.56 | ~ 12-14 | Slow | (16) |
| Q151M | 2 | Tv | 0.02 | 2.95 | 0.03 | 45.45 | ~ 15-18 | Slow | (26) |
| L210W | 1 | Tv | 0.56 | 27.5 | 0.57 | 49.50 | ~ 15-24 | Slow | (25) |

**Table S3.** Acquisition times of emergence of major non-nucleoside RT inhibitor (NNRTI) resistance mutations using KMC simulation and literature survey, which can occur as a transition (Ti), transversion (Tv) or both, and involve different numbers of nucleotide changes (Δnuc). The table is ordered by the measured by the estimated time constant **τ**.

| Mutations | $\Delta\text{nuc}$ | Nature of Mutation | Frequency (%) | | | Time Constant ($\tau$) | Timeline in Literature (Months) | Classification of Timeline (Literature) | Reference |
| --- | --- | --- | --- | --- | --- | --- | --- | --- | --- |
| | | | Drug-naive | Drug-exper. | Adaptive Frequency $\langle F_a^l(t=0) \rangle$ | | | | |
| V106M | 2 | Ti | 0.04 | 1 | 0.62 | 4.80 | ~ 0.25-0.5 | Fast | (27) |
| K103N | 1 | Tv | 2.75 | 36 | 26.0 | 5.58 | ~ 0.25-1 | Fast | (28, 29) |
| E138K | 1 | Ti | 0.16 | 1 | 0.67 | 7.88 | ~ 0.3-2.7 | Fast | (29, 30) |
| V106A | 1 | Ti | 0.04 | 1 | 1.2 | 9.54 | ~ 1-3 | Fast | (29, 31) |
| Y188L | 2 | Tv | 0.21 | 5 | 3.2 | 17.18 | ~ 3-4 | Intermed. | (32) |
| Y181V | 2 | Tv | 0.008 | 1 | 0.18 | 20.31 | ~ 4-12 | Intermed.-Slow | (33-35) |
| L100I | 1 | Tv | 0.09 | 5 | 0.66 | 23.18 | ~ 5-7 | Intermed.-Slow | (36) |
| Y181C | 1 | Ti | 0.44 | 15.2 | 5.5 | 25.15 | ~ 6-12 | Slow | (34, 37) |
| Y181I | 2 | Tv | 0.01 | 1 | 0.38 | 26.59 | ~ 6-12 | Slow | (33) |
| K101E | 1 | Ti | 0.28 | 7.3 | 1.8 | 26.60 | ~ 7-8 | Slow | (36) |
| G190A | 1 | Tv | 0.36 | 13.6 | 1.7 | 26.66 | ~6-12 | Slow | (38) |

**Table S4.** Acquisition times of emergence of major INSTI resistance mutations using KMC simulation and literature survey, which can occur as a transition (Ti), transversion (Tv) or both, and involve different numbers of nucleotide changes (Δnuc). The table is ordered by the measured by the estimated time constant **τ**.

| Mutations | $\Delta\text{nuc}$ | Nature of Mutation | Frequency (%) | | | Time Constant ($\tau$) | Timeline in Literature (Months) | Classification of Timeline (Literature) | Reference |
| --- | --- | --- | --- | --- | --- | --- | --- | --- | --- |
| | | | Drug-Naive | Drug-Experienced | Adaptive Frequency $\langle F_a^l(t=0) \rangle$ | | | | |
| E92Q | 1 | Tv | 0.02 | 3.2 | 2.2 | 2.1 | ~ 0.25 | Fast | (39) |
| N155H | 1 | Tv | 0 | 24.3 | 22.3 | 2.9 | ~0.25-0.5 | Fast | (39, 40) |
| Y143R | 2 | Ti | 0.05 | 5.5 | 2.6 | 3.8 | ~ 3 | Fast | (41) |
| V151I | 1 | Ti | 0.03 | 11.3 | 5.8 | 6.6 | ~ 1-5 | Fast | (42, 43) |
| Q148R | 1 | Ti | 0.06 | 5.0 | 1.9 | 21.5 | ~ 3-4 | Intermed. | (44) |
| Y143C | 1 | Ti | 0.10 | 2.4 | 0.15 | 27.6 | ~ 6-12 | Slow | (45) |
| S230R | 1 | Tv | 0.07 | 24.6 | 0.12 | 34.2 | ~ 7 | Slow | (46) |
| G140A | 1 | Tv | 0.42 | 1.1 | 0.8 | 42.0 | ~ 6-8 | Slow | (47) |
| E138A | 1 | Tv | 0.0 | 1.7 | 0.22 | 43.3 | ~ 6-12 | Slow | (47) |
| S147G | 2 | Tv | 0.02 | 1.0 | 0.20 | 45.9 | ~ 6 | Slow | (48) |
| Q148K | 1 | Tv | 0.10 | 1.0 | 0.12 | 49.5 | ~ 12 | Slow | (49) |
| G140S | 1 | Ti | 0.12 | 19.9 | 0.23 | 64.1 | ~ 12 – 16 | Slow | (50-52) |
| Q148H | 1 | Tv | 0.02 | 18.6 | 0.95 | 64.9 | ~ 12 – 16 | Slow | (41, 51, 52) |
| E138K | 1 | Ti | 1.2 | 3.4 | 0.29 | 86.9 | ~ 7-16 | Slow | (47) |

**Figure S4.**
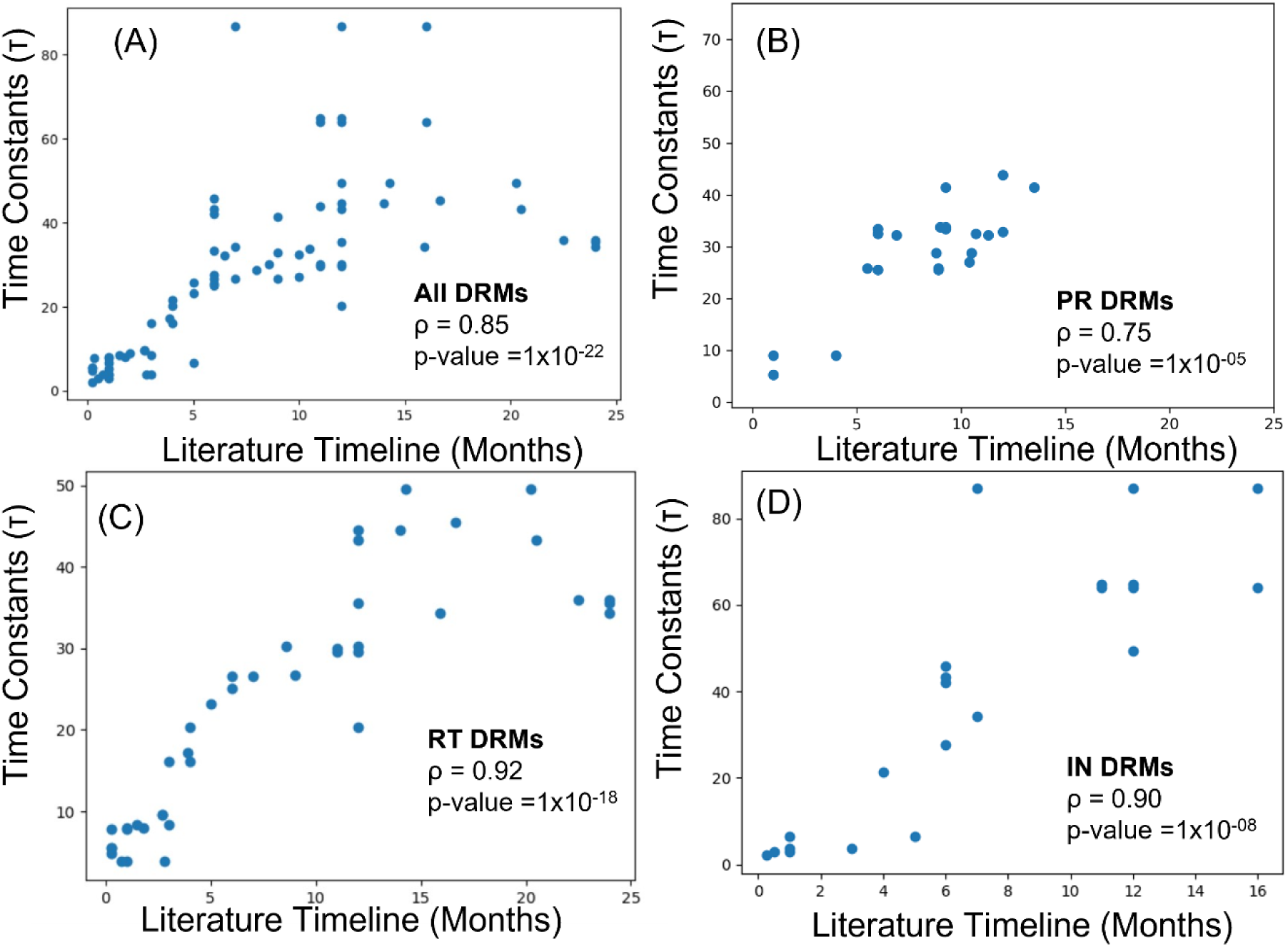
Spearman rank correlation between the literature timeline for DRMs to appear in drug-experienced population and the Potts+KMC simulation predicted time constants (τ) in (A) All mutations, (B) protease (PR) (C) reverse transcriptase (RT/NRTI+NNRTI) and (D) integrase (IN).

**Figure S5:**
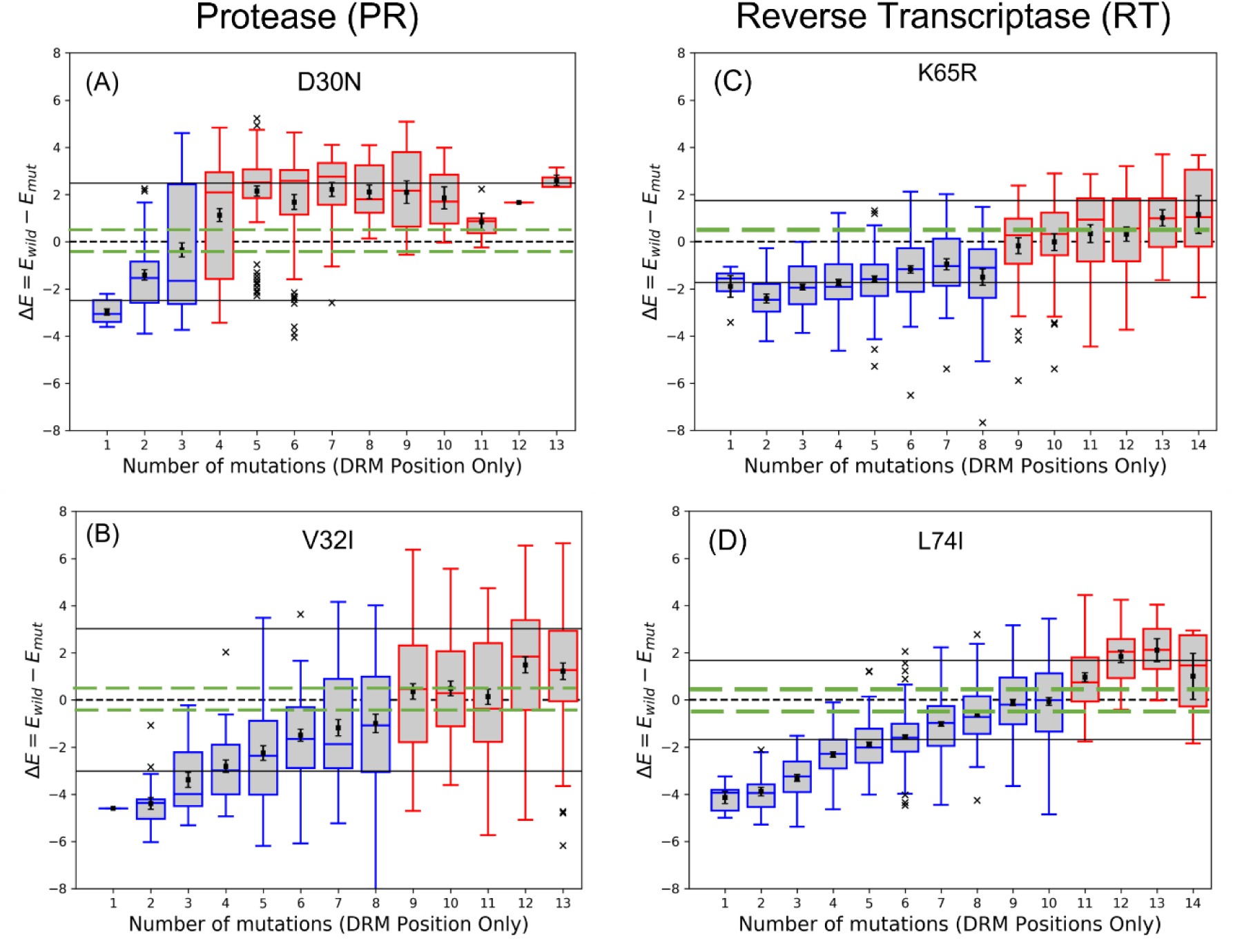
The effect of epistasis on the favorability of a primary resistance mutation in protease (PR) and reverse transcriptase (RT). Change of Potts energy difference (ΔE) for a mutation of sequences with the particular mutation, conditional on the total number mutations with respect to wild type consensus considering only the drug resistance positions (∼36 for PR and ∼31 for RT) is shown as boxplots annotating the first, second, and third interquartile range. The whiskers extend to 1.5 times the interquartile range with outliers marked as ’x’s and the mean values are marked as squares. The left ordinate scale shows the Potts energy difference (ΔE). The area around ΔE = 0 indicated with green dashed lines (−0.5 < ΔE < 0.5) defines the neutral zone. The sequences whose energy difference falls above ΔE = 0.5 (short-dashed line) are entrenching backgrounds favoring the mutation. Sequence backgrounds where the mutation is favored on average are shown in red box plots, the others in blue box plots. (A) The mutation D30N with faster kinetics becomes favorable on average when there are more than ∼4 total mutations at the drug resistance positions with respect to wild type consensus. (B) The mutation V32I with slower kinetics becomes favorable on average when there are more than ∼9 total mutations at the drug resistance positions with respect to wild type consensus. (C) The mutation K65R with faster kinetics becomes favorable on average when there are more than ∼9 total mutations at the drug resistance positions with respect to wild type consensus. (D) The mutation L74I with slower kinetics becomes favorable on average when there are more than ∼11 total number of mutations at the drug resistance positions with respect to wild type consensus. The black dashed line represents ΔE = 0. The black solid lines represent the standard deviation for ΔE. The black dashed line represents ΔE = 0. The black solid lines represent the standard deviation for ΔE.

**Figure S6:**
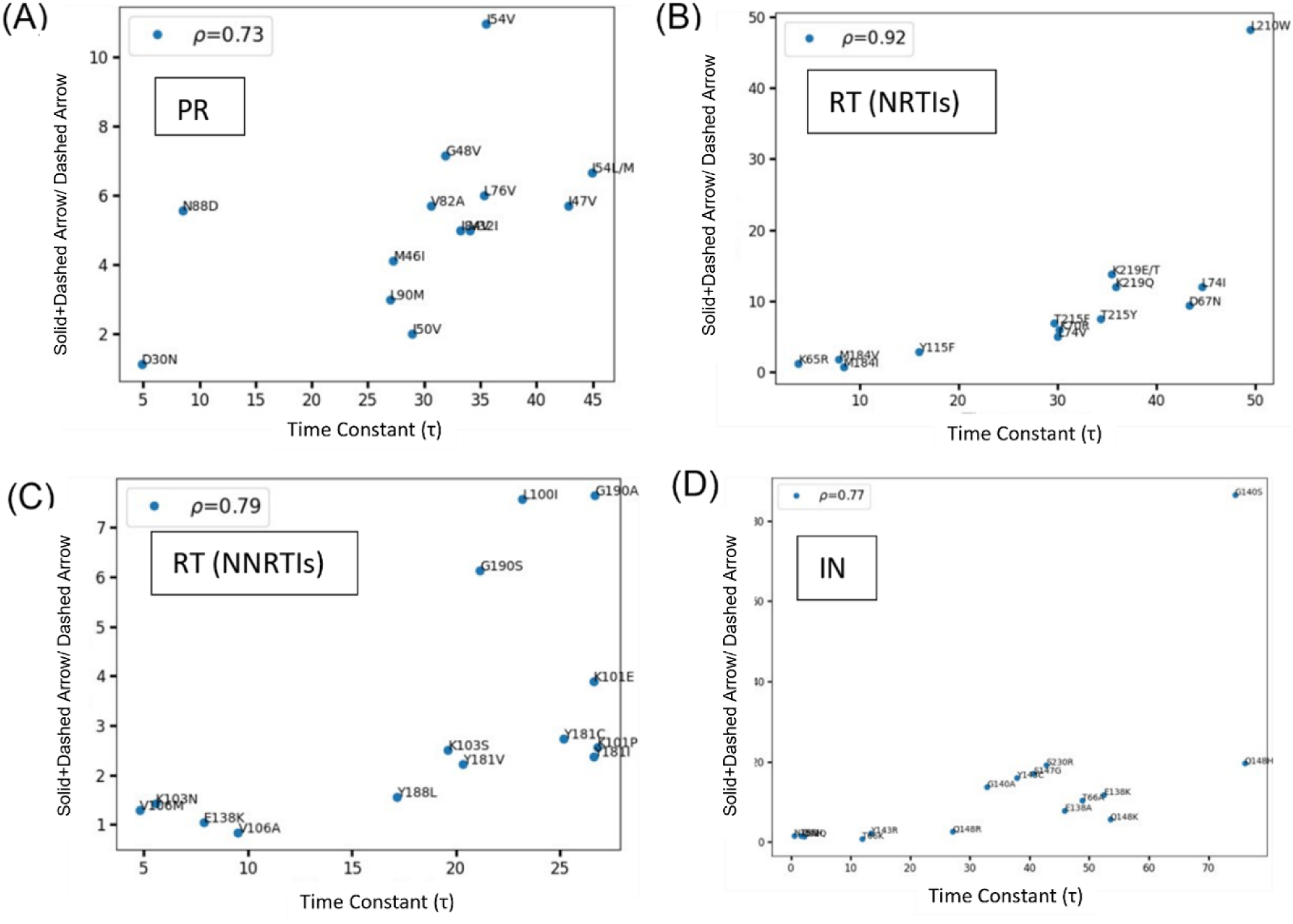
Spearman rank correlation between the new drug-exposed environment (sum of solid + dashed arrow from Fig. 3) to the initial increase in adaptive frequency (size of the dashed arrow) and the Potts+KMC simulation predicted time constants (τ) in different protein targets such as (A) protease (PR), (B) reverse transcriptase (RT/NRTI), (C) reverse transcriptase (RT/NNRTI) and (D) integrase (IN).

**Figure S7:**
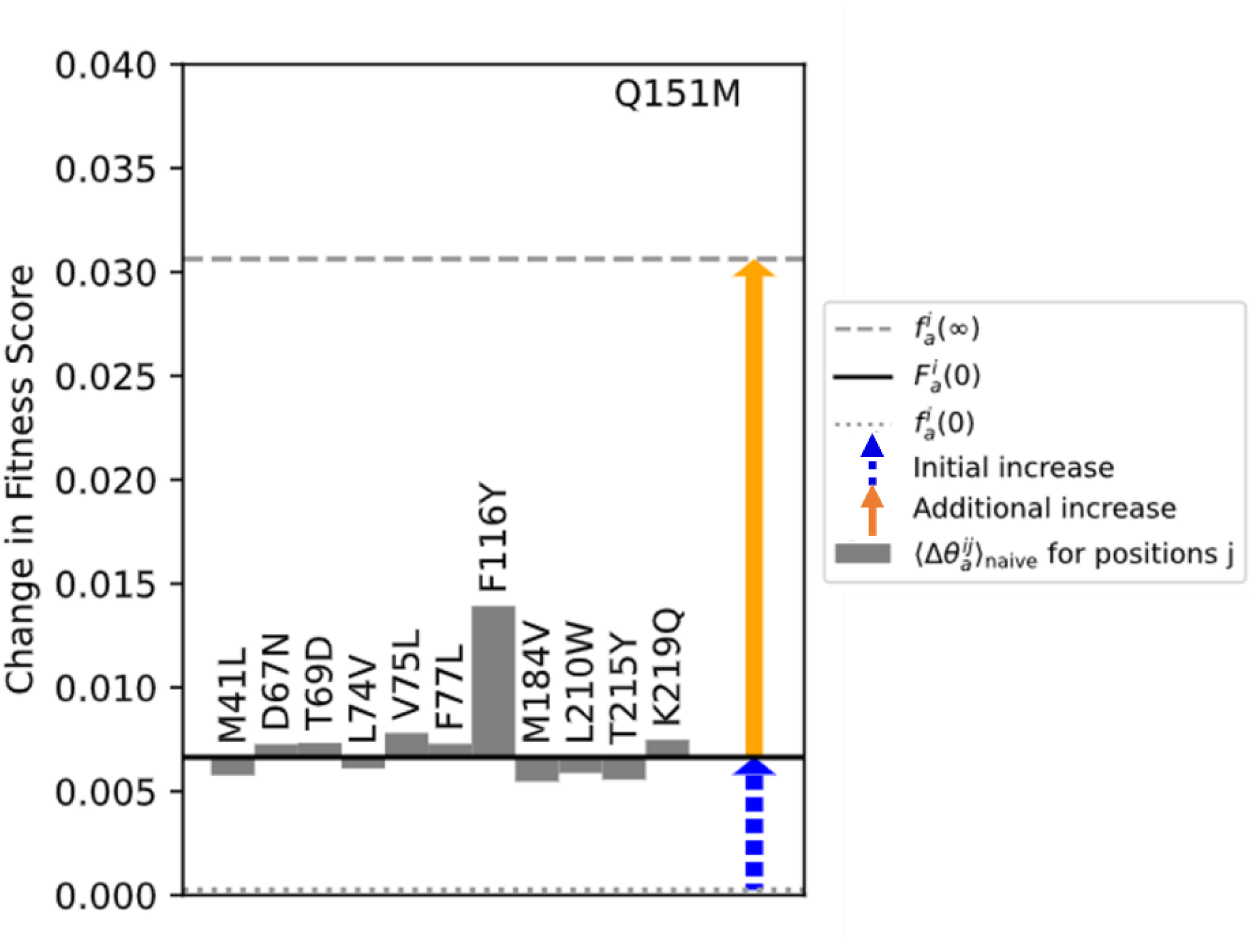
Comparison of the additional mutations necessary for Q151M (RT/NRTI) to become favored. In this plot, the focal DRM (Q151M) is listed above the dashed blue and solid orange arrows. The dotted grey line reflects the DRM’s frequency in the drug-naïve ensemble, the solid black line reflects the DRM’s average fitness under the drug-experienced Hamiltonian in the drug-naïve ensemble, and the dashed grey line reflects the DRM’s frequency in the drug-experienced ensemble. The dashed blue and solid orange arrows are identical to the arrows in **Fig. 3 and 4** (Main text). The dashed blue arrow then represents the initial increase in the DRM’s frequency upon exposure to drugs starting from the drug-naïve state if the background were held fixed, identical to the dashed blue arrows in **Fig. 3 and 4**, and the solid orange arrow represents the additional increase in DRM frequency once the sequence backgrounds are allowed to vary and accumulate coupled mutations, identical to the solid orange arrows in **Fig. 3** (Main text). The grey bars represent the fitness measured using equation 2 (Main text), averaged over the drug-naïve ensemble, giving the first-order contribution to the increase in the fitness of the DRM at i due to the evolution of j to the drug-experienced state.. The sum of the grey bar magnitudes closely approximates the length of the solid orange arrow.

### Section S1

#### Literature evidence

The impact of identified compensatory mutations on the kinetics of "fast" and "slow" drug resistance mutations.

In this section, we will examine literature evidence demonstrating the co-occurrence of focal "slow" and "fast" drug resistance mutations (DRMs) alongside their corresponding background mutations for each respective enzymatic drug target, as depicted in Fig. 4 in the main text. In PR, the “fast” and “slow” focal DRMs are D30N and V32I, respectively. Based on existing literature (53, 54), we know that the nonpolymorphic accessory mutation N88D is frequently selected by nelfinavir (NFV) in conjunction with D30N. Notably, Sugiura et al. (55) reported a substantial suppression in the acquisition of D30N in the presence of L90M; in support of this, we also observe a negative biasing effect of L90M on the D30N focal DRM (**Fig. 4A** **(left)**). Thus, our study validates these experimental observations, as depicted in (**Fig. 4A** **(right)**), by demonstrating that the "slow" mutation V32I exhibits a change in adaptive frequency influenced by background mutational patterns, namely M46I, I47V, A71V, and V82A, which collectively act as an epistatic barrier. In nearly two-thirds of the sequences, V32I is observed in combination with I47V/A, resulting in high-level resistance to lopinavir (LPV) and intermediate resistance to darunavir (DRV), as reported in the Stanford database (56). Furthermore, other literature sources confirm the co-occurrence of V32I with other drug resistance mutations (DRMs), such as M46I and V82A, in response to saquinavir (SQV), LPV, and DRV (57). Similar evidence of concurrent background mutation patterns, as identified in our study (**Fig. 4B-C**), alongside our focal "slow" DRMs, can be found in the literature for the target enzyme RT (NRTIs and NNRTIs). Among these, M184V (fast DRM) and D67N (slow DRM) are significant NRTI resistance mutations. D67N serves as a prototypical example of a non-polymorphic mutation selected by thymidine analogs zidovudine (AZT) and stavudine (d4T), known as thymidine analog mutations or TAMs (1). Specifically, D67N falls under the type 2 TAMs mutational patterns and frequently co-occurs with other TAM-associated DRMs, including K70R, T215F/Y, and K219Q/E (58-60). Conversely, M184V is selected by lamivudine/emtricitabine (3TC/FTC) and leads to a >200-fold reduction in susceptibility to these drugs (61, 62) and is commonly detected in patients experiencing virological failure on 3TC or FTC (63, 64). The most prevalent pattern of mutations associated with patients receiving abacavir/lamivudine (ABC/3TC) involves the combination of L74V and M184V, which reduces abacavir (ABC) susceptibility by approximately 5-fold (65-67). In **Fig. 4C**, we illustrate K103N (fast DRM) and Y181C (slow DRM) as examples of NNRTI mutations. According to the literature, K103N often occurs in conjunction with L100I (68), V179I/D/E (69), and P225H (58), which we also identify. Similarly, Y181C has been found to co-occur with mutations such as M184V, T215Y, and K219Q (70), G190A (71), and M221V (72), consistent with our prediction of compensatory mutations in the background. Another slowly evolving NRTI DRM, Q151M (**Fig. S7**) is predominantly selected by AZT and ddI, the first available NRTI drugs and emerges after (∼15 months) (73). Typically, it manifests alongside two or more of the subsequent four accessory mutations: A62V, V75I, F77L, and F116Y (73-75); this so-called “Q151M complex” mutant (A62V/V75I/F77L/F116Y/Q151M) is referred to as Q151Mc (73-75). **Fig. S7** predicts three of these accessory mutations (V75I, F77L, and particularly F116Y) established in the literature. The structural analysis of the Q151M complex (slow RT/NRTI DRM, **Fig. S7**) in various nucleic-acid bound states of HIV-1 RT suggests that the F116Y mutation leads to formation of hydrogen bond of the phenoxyl ring with the main-chain carbonyl group of Lys73, which results in restricting the conformational flexibility of the M151 side chain and helping to compensate for the loss in RT activity, and hence fitness, that is experienced with the Q151M mutation alone (76). In **Fig. 4D**, we present a pair of INSTI mutations, N155H (fast mutation) and Q148H (slow mutation), along with their respective list of predicted compensatory mutations. Despite having very different timescales for onset, these two mutations are some of the most frequently encountered DRMs under INSTI selection pressure (77). The fast mutation N155H arises in combination with numerous other background DRMs, but Q148H is most commonly found in combination with G140S. Indeed, our prediction confirms the numerous literature reports (78-80) for the significant contribution of G140S to the adaptive frequency of Q148H. The N155H mutants emerge first, and are eventually replaced by Q148H and Y143R mutants (81), usually in combination with G140S (82) which explains the decrease in N155H frequency (downward black solid arrow in **Fig. 4D**) with time compare to its adaptive-frequency.

## References

1. UNAIDS (2022) Global HIV & AIDS statistics — Fact sheet. URL: https://www.unaids.org/en/resources/fact-sheet. (UNAIDS).

2. J. M. Coffin, The virology of AIDS: 1990. AIDS 4, S9 (1990).

3. J. M. Coffin, HIV Population Dynamics in Vivo: Implications for Genetic Variation, Pathogenesis, and Therapy. Science 267, 483–489 (1995).

4. A. S. Perelson, A. U. Neumann, M. Markowitz, J. M. Leonard, D. D. Ho, HIV-1 Dynamics in Vivo: Virion Clearance Rate, Infected Cell Life-Span, and Viral Generation Time. Science 271, 1582–1586 (1996).

5. S.-Y. Rhee et al., Public availability of HIV-1 drug resistance sequence and treatment data: a systematic review. The Lancet Microbe 3, e392–e398 (2022).

6. J. M. Cuevas, R. Geller, R. Garijo, J. López-Aldeguer, R. Sanjuán, Extremely High Mutation Rate of HIV-1 In Vivo. PLoS Biol. 13, e1002251 (2015).

7. J. P. Barton et al., Relative rate and location of intra-host HIV evolution to evade cellular immunity are predictable. Nat. Commun. 7, 11660 (2016).

8. P. L. Boyer, S. G. Sarafianos, E. Arnold, S. H. Hughes, Selective excision of AZTMP by drug-resistant human immunodeficiency virus reverse transcriptase. J Virol 75, 4832–4842 (2001).

9. S. G. Sarafianos et al., Structure and function of HIV-1 reverse transcriptase: molecular mechanisms of polymerization and inhibition. J. Mol. Biol. 385, 693–713 (2009).

10. D. O. Passos et al., Structural basis for strand-transfer inhibitor binding to HIV intasomes. Science 367, 810–814 (2020).

11. M. F. Kearney et al., Lack of detectable HIV-1 molecular evolution during suppressive antiretroviral therapy. PLoS Pathog. 10, e1004010 (2014).

12. J. I. Boucher et al., Constrained Mutational Sampling of Amino Acids in HIV-1 Protease Evolution. Mol. Biol. Evol. 36, 798–810 (2019).

13. E. Spielvogel et al., Selection of HIV-1 for resistance to fifth-generation protease inhibitors reveals two independent pathways to high-level resistance. eLife 12 (2023).

14. T. C. Butler, J. P. Barton, M. Kardar, A. K. Chakraborty, Identification of drug resistance mutations in HIV from constraints on natural evolution. *Phys*. Rev. E 93, 022412 (2016).

15. S. W. Lockless, R. Ranganathan, Evolutionarily Conserved Pathways of Energetic Connectivity in Protein Families. Science 286, 295–299 (1999).

16. J. D. Bloom, L. I. Gong, D. Baltimore, Permissive Secondary Mutations Enable the Evolution of Influenza Oseltamivir Resistance. Science 328, 1272–1275 (2010).

17. O. Haq, M. Andrec, A. V. Morozov, R. M. Levy, Correlated Electrostatic Mutations Provide a Reservoir of Stability in HIV Protease. PLoS Comput. Biol. 8, e1002675 (2012).

18. R. M. Levy, A. Haldane, W. F. Flynn, Potts Hamiltonian models of protein co-variation, free energy landscapes, and evolutionary fitness. Curr Opin Struct Biol 43, 55–62 (2017).

19. P. Tian, J. M. Louis, J. L. Baber, A. Aniana, R. B. Best, Co-Evolutionary Fitness Landscapes for Sequence Design. Angew Chem. Int. Ed. Engl. 57, 5674-5678 (2018).

20. R. A. Neher, B. I. Shraiman, Competition between recombination and epistasis can cause a transition from allele to genotype selection. Proc Natl Acad Sci U S A 106, 6866–6871 (2009).

21. M. W. Chang, B. E. Torbett, Accessory Mutations Maintain Stability in Drug-Resistant HIV-1 Protease. J. Mol. Biol. 410, 756–760 (2011).

22. W. F. Flynn et al., Deep Sequencing of Protease Inhibitor Resistant HIV Patient Isolates Reveals Patterns of Correlated Mutations in Gag and Protease. PLoS Comput. Biol. 11, e1004249 (2015).

23. Stanford (2023) Stanford HIV Database. URL: https://hivdb.stanford.edu/. (Stanford).

24. L. Alamos (2023) Los Alamos HIV sequence database. URL: http://www.hiv.lanl.gov/.

25. A. M. Wensing et al., 2022 update of the drug resistance mutations in HIV-1. Top Antivir. Med. 30, 559–574 (2022).

26. A. L. Ferguson et al., Translating HIV sequences into quantitative fitness landscapes predicts viral vulnerabilities for rational immunogen design. Immunity 38, 606–617 (2013).

27. J. P. Barton, M. Kardar, A. K. Chakraborty, Scaling laws describe memories of host-pathogen riposte in the HIV population. Proc. Natl. Acad. Sci. U.S.A. 112, 1965–1970 (2015).

28. D. D. Pollock, G. Thiltgen, R. A. Goldstein, Amino acid coevolution induces an evolutionary Stokes shift. Proc. Natl. Acad. Sci. U.S.A. 109, E1352–E1359 (2012).

29. P. Shah, D. M. McCandlish, J. B. Plotkin, Contingency and entrenchment in protein evolution under purifying selection. Proc. Natl. Acad. Sci. U.S.A. 112, E3226–E3235 (2015).

30. D. M. McCandlish, P. Shah, J. B. Plotkin, Epistasis and the Dynamics of Reversion in Molecular Evolution. Genetics 203, 1335–1351 (2016).

31. M. A. Depristo, D. M. Weinreich, D. L. Hartl, Missense meanderings in sequence space: a biophysical view of protein evolution. Nat. Rev. Genet. 6, 678–687 (2005).

32. M. J. Harms, J. W. Thornton, Historical contingency and its biophysical basis in glucocorticoid receptor evolution. Nature 512, 203–207 (2014).

33. N. K. Yilmaz, C. A. Schiffer, "Drug Resistance to HIV-1 Protease Inhibitors: Molecular Mechanisms and Substrate Coevolution". (Springer International Publishing, 2017), 10.1007/978-3-319-46718-4_35, pp. 535-544.

34. A. Biswas, A. Haldane, E. Arnold, R. M. Levy, Epistasis and entrenchment of drug resistance in HIV-1 subtype B. Elife 8 (2019).

35. Y. Iwasa, F. Michor, M. A. Nowak, Stochastic tunnels in evolutionary dynamics. Genetics 166, 1571–1579 (2004).

36. Y. Guo, M. Vucelja, A. Amir, Stochastic tunneling across fitness valleys can give rise to a logarithmic long-term fitness trajectory. Sci. Adv. 5, eaav3842 (2019).

37. I. Choudhuri, A. Biswas, A. Haldane, R. M. Levy, Contingency and Entrenchment of Drug-Resistance Mutations in HIV Viral Proteins. J. Phys. Chem. B (2022).

38. A. Biswas, A. Haldane, R. M. Levy, Limits to detecting epistasis in the fitness landscape of HIV. PLoS One 17, e0262314 (2022).

39. J. A. de la Paz, C. M. Nartey, M. Yuvaraj, F. Morcos, Epistatic contributions promote the unification of incompatible models of neutral molecular evolution. Proc. Natl. Acad. Sci. U.S.A. 117, 5873–5882 (2020).

40. J. Gizzio, A. Thakur, A. Haldane, R. M. Levy, Evolutionary divergence in the conformational landscapes of tyrosine vs serine/threonine kinases. Elife 11 (2022).

41. A. Haldane, R. M. Levy, Mi3-GPU: MCMC-based inverse Ising inference on GPUs for protein covariation analysis. Comput. Phys. Commun. 260, 107312 (2021).

42. A. Chaillon et al., HIV persists throughout deep tissues with repopulation from multiple anatomical sources. J. Clin. Invest. 130, 1699–1712 (2020).

43. K. Theys et al., The impact of HIV-1 within-host evolution on transmission dynamics. Curr. Opin. Virol. 28, 92–101 (2018).

44. A. F. Feder, K. N. Harper, C. J. Brumme, P. S. Pennings, Understanding patterns of HIV multi-drug resistance through models of temporal and spatial drug heterogeneity. Elife 10 (2021).

45. A. R. Wargo, G. Kurath, Viral fitness: definitions, measurement, and current insights. Curr. Opin. Virol. 2, 538–545 (2012).

46. E. Domingo, J. J. Holland, RNA virus mutations and fitness for survival. Annu. Rev. Microbiol. 51, 151–178 (1997).

47. G. Sella, A. E. Hirsh, The application of statistical physics to evolutionary biology. Proc Natl Acad Sci U S A 102, 9541–9546 (2005).

48. S. C. Bihani, G. D. Gupta, M. V. Hosur, Molecular basis for reduced cleavage activity and drug resistance in D30N HIV-1 protease. J. Biomol. Struct. Dyn. 40, 13127–13135 (2022).

49. S. Pawar et al., Structural studies of antiviral inhibitor with HIV-1 protease bearing drug resistant substitutions of V32I, I47V and V82I. Biochem. Biophys. Res. Commun. 514, 974-978 (2019).

50. D. A. Ragland et al., Drug resistance conferred by mutations outside the active site through alterations in the dynamic and structural ensemble of HIV-1 protease. J. Am. Chem. Soc. 136, 11956–11963 (2014).

51. X. Tu et al., Structural basis of HIV-1 resistance to AZT by excision. Nat. Struct. Mol. Biol. 17, 1202–1209 (2010).

52. M. T. Lai et al., Mechanistic Study of Common Non-Nucleoside Reverse Transcriptase Inhibitor-Resistant Mutations with K103N and Y181C Substitutions. Viruses 8 (2016).

53. Y. Hsiou et al., The Lys103Asn mutation of HIV-1 RT: a novel mechanism of drug resistance. J. Mol. Biol. 309, 437–445 (2001).

54. K. Das et al., Crystal structures of 8-Cl and 9-Cl TIBO complexed with wild-type HIV-1 RT and 8-Cl TIBO complexed with the Tyr181Cys HIV-1 RT drug-resistant mutant. J. Mol. Biol. 264, 1085–1100 (1996).

55. J. Ren et al., Structural mechanisms of drug resistance for mutations at codons 181 and 188 in HIV-1 reverse transcriptase and the improved resilience of second generation non-nucleoside inhibitors. J. Mol. Biol. 312, 795–805 (2001).

56. S. Hare et al., Molecular mechanisms of retroviral integrase inhibition and the evolution of viral resistance. Proc. Natl. Acad. Sci. U.S.A. 107, 20057–20062 (2010).

57. N. J. Cook et al., Structural basis of second-generation HIV integrase inhibitor action and viral resistance. Science 367, 806–810 (2020).

58. M. Li et al., Mechanisms of HIV-1 integrase resistance to dolutegravir and potent inhibition of drug-resistant variants. Sci. Adv. 9, eadg5953 (2023).

59. P. L. Tzou et al., Integrase strand transfer inhibitor (INSTI)-resistance mutations for the surveillance of transmitted HIV-1 drug resistance. J. Antimicrob. Chemother. 75, 170–182 (2020).

60. W. F. Flynn, A. Haldane, B. E. Torbett, R. M. Levy, Inference of Epistatic Effects Leading to Entrenchment and Drug Resistance in HIV-1 Protease. Mol. Biol. Evol. 34, 1291–1306 (2017).

61. E. F. Pettersen et al., UCSF Chimera--a visualization system for exploratory research and analysis. J. Comput. Chem. 25, 1605–1612 (2004).

62. S. Cocco, C. Feinauer, M. Figliuzzi, R. Monasson, M. Weigt, Inverse statistical physics of protein sequences: a key issues review. Rep. Prog. Phys. 81, 032601 (2018).

63. M. Mézard, T. Mora, Constraint satisfaction problems and neural networks: A statistical physics perspective. J. Physiol. Paris. 103, 107–113 (2009).

64. M. Weigt, R. A. White, H. Szurmant, J. A. Hoch, T. Hwa, Identification of direct residue contacts in protein-protein interaction by message passing. Proc. Natl. Acad. Sci. U.S.A. 106, 67–72 (2009).

65. J. I. Sułkowska, F. Morcos, M. Weigt, T. Hwa, J. N. Onuchic, Genomics-aided structure prediction. Proc. Natl. Acad. Sci. U.S.A. 109, 10340–10345 (2012).

66. R. Nassar, G. L. Dignon, R. M. Razban, K. A. Dill, The Protein Folding Problem: The Role of Theory. J. Mol. Biol. 433, 167126 (2021).

67. F. Morcos, N. P. Schafer, R. R. Cheng, J. N. Onuchic, P. G. Wolynes, Coevolutionary information, protein folding landscapes, and the thermodynamics of natural selection. Proc. Natl. Acad. Sci. U.S.A. 111, 12408–12413 (2014).

68. N. Bhattacharya et al., Interpreting Potts and Transformer Protein Models Through the Lens of Simplified Attention. Pac. Symp. Biocomput. 27, 34–45 (2022).

69. W. K. Hastings, Monte Carlo Sampling Methods Using Markov Chains and Their Applications. Biometrika 57, 97 (1970).

## Reference

1. Stanford (2023) Stanford HIV Database. URL: https://hivdb.stanford.edu/. (Stanford).

2. M. Markowitz et al., A Preliminary Evaluation of Nelfinavir Mesylate, an Inhibitor of Human Immunodeficiency Virus (HIV)-1 Protease, to Treat HIV Infection. J. Infect. Dis. 177, 1533-1540 (1998).

3. D. J. Kempf et al., Incidence of resistance in a double-blind study comparing lopinavir/ritonavir plus stavudine and lamivudine to nelfinavir plus stavudine and lamivudine. J. Infect. Dis. 189, 51-60 (2004).

4. J. E. Fitzgibbon et al., Emergence of Drug Resistance Mutations in a Group of HIV-Infected Children Taking Nelfinavir-Containing Regimens. AIDS Res. Hum. Retrovir. 17, 1321-1328 (2001).

5. J. H. Condra et al., Genetic correlates of in vivo viral resistance to indinavir, a human immunodeficiency virus type 1 protease inhibitor. Journal of Virology 70, 8270-8276 (1996).

6. J. Lawrence et al., Clinical resistance patterns and responses to two sequential protease inhibitor regimens in saquinavir and reverse transcriptase inhibitor-experienced persons. J. Infect. Dis. 179, 1356-1364 (1999).

7. T. J. Barber et al., Frequency and patterns of protease gene resistance mutations in HIV-infected patients treated with lopinavir/ritonavir as their first protease inhibitor. J. Antimicrob. Chemother. 67, 995-1000 (2012).

8. G. Sterrantino et al., Genotypic resistance profiles associated with virological failure to darunavir-containing regimens: a cross-sectional analysis. Infection 40, 311-318 (2012).

9. Y. M. Zhang et al., Drug resistance during indinavir therapy is caused by mutations in the protease gene and in its Gag substrate cleavage sites. J. Virol. 71, 6662-6670 (1997).

10. D. Wang et al., Evolution of drug-resistant viral populations during interruption of antiretroviral therapy. J. Virol. 85, 6403-6415 (2011).

11. R. Kantor et al., Evolution of Primary Protease Inhibitor Resistance Mutations during Protease Inhibitor Salvage Therapy. Antimicrob. Agents Chemother. 46, 1086-1092 (2002).

12. S. de Meyer et al., Resistance profile of darunavir: combined 24-week results from the POWER trials. AIDS Res. Hum. Retroviruses 24, 379-388 (2008).

13. L. Bélec et al., High levels of drug-resistant human immunodeficiency virus variants in patients exhibiting increasing CD4+ T cell counts despite virologic failure of protease inhibitor-containing antiretroviral combination therapy. J. Infect. Dis. 181, 1808-1812 (2000).

14. N. Shulman et al., Virtual inhibitory quotient predicts response to ritonavir boosting of indinavir-based therapy in human immunodeficiency virus-infected patients with ongoing viremia. Antimicrob. Agents Chemother. 46, 3907-3916 (2002).

15. J. F. Delfraissy et al., Lopinavir/ritonavir monotherapy or plus zidovudine and lamivudine in antiretroviral-naive HIV-infected patients. Aids 22, 385-393 (2008).

16. M. Wirden et al., Antiretroviral combinations implicated in emergence of the L74I and L74V resistance mutations in HIV-1-infected patients. AIDS 23 (2009).

17. M. Maguire et al., Emergence of resistance to protease inhibitor amprenavir in human immunodeficiency virus type 1-infected patients: selection of four alternative viral protease genotypes and influence of viral susceptibility to coadministered reverse transcriptase nucleoside inhibitors. Antimicrob. Agents Chemother. 46, 731-738 (2002).

18. J. G. GarcíA-Lerma et al., A Novel Genetic Pathway of Human Immunodeficiency Virus Type 1 Resistance to Stavudine Mediated by the K65R Mutation. J. Virol. 77, 5685-5693 (2003).

19. P. J. Ruane, A. D. Luber, K65R-associated virologic failure in HIV-infected patients receiving tenofovir-containing triple nucleoside/nucleotide reverse transcriptase inhibitor regimens. MedGenMed 6, 31 (2004).

20. M. Tisdale, T. Alnadaf, D. Cousens, Combination of mutations in human immunodeficiency virus type 1 reverse transcriptase required for resistance to the carbocyclic nucleoside 1592U89. Antimicrob. Agents Chemother. 41, 1094-1098 (1997).

21. N. A. Margot, J. M. Waters, M. D. Miller, In Vitro Human Immunodeficiency Virus Type 1 Resistance Selections with Combinations of Tenofovir and Emtricitabine or Abacavir and Lamivudine. Antimicrob. Agents Chemother. 50, 4087-4095 (2006).

22. D. J. Hooker et al., An in vivo mutation from leucine to tryptophan at position 210 in human immunodeficiency virus type 1 reverse transcriptase contributes to high-level resistance to 3’-azido-3’-deoxythymidine. J. Virol. 70, 8010-8018 (1996).

23. S. Yerly et al., Switch to Unusual Amino Acids at Codon 215 of the Human Immunodeficiency Virus Type 1 Reverse Transcriptase Gene in Seroconvertors Infected with Zidovudine-Resistant Variants. J. Virol. 72, 3520-3523 (1998).

24. M. D. Miller et al., Genotypic and Phenotypic Predictors of the Magnitude of Response to Tenofovir Disoproxil Fumarate Treatment in Antiretroviral-Experienced Patients. J. Infect. Dis. 189, 837-846 (2004).

25. N. Yahi, C. Tamalet, C. Tourrès, N. Tivoli, J. Fantini, Mutation L210W of HIV-1 reverse transcriptase in patients receiving combination therapy. J. Biomed. Sci. 7, 507-513 (2000).

26. S. T. Burda et al., HIV-1 reverse transcriptase drug-resistance mutations in chronically infected individuals receiving or naive to HAART in Cameroon. J. Med. Virol. 82, 187-196 (2010).

27. M. Feng et al., *In Vitro* Resistance Selection with Doravirine (MK-1439), a Novel Nonnucleoside Reverse Transcriptase Inhibitor with Distinct Mutation Development Pathways. Antimicrob. Agents Chemother. 59, 590-598 (2015).

28. L. T. Bacheler et al., Human Immunodeficiency Virus Type 1 Mutations Selected in Patients Failing Efavirenz Combination Therapy. Antimicrob. Agents Chemother. 44, 2475-2484 (2000).

29. M. Feng et al., In vitro resistance selection with doravirine (MK-1439), a novel nonnucleoside reverse transcriptase inhibitor with distinct mutation development pathways. Antimicrob. Agents Chemother. 59, 590-598 (2015).

30. R. Kulkarni et al., The HIV-1 Reverse Transcriptase M184I Mutation Enhances the E138K-Associated Resistance to Rilpivirine and Decreases Viral Fitness. J. Acquir. Immune Defic. Syndr. 59 (2012).

31. J. Balzarini, H. Pelemans, R. Esnouf, E. De Clercq, A novel mutation (F227L) arises in the reverse transcriptase of human immunodeficiency virus type 1 on dose-escalating treatment of HIV type 1-infected cell cultures with the nonnucleoside reverse transcriptase inhibitor thiocarboxanilide UC-781. AIDS Res. Hum. Retroviruses 14, 255-260 (1998).

32. L. Bacheler et al., Genotypic Correlates of Phenotypic Resistance to Efavirenz in Virus Isolates from Patients Failing Nonnucleoside Reverse Transcriptase Inhibitor Therapy. J. Virol. 75, 4999-5008 (2001).

33. L. Tambuyzer et al., Characterization of genotypic and phenotypic changes in HIV-1-infected patients with virologic failure on an etravirine-containing regimen in the DUET-1 and DUET-2 clinical studies.

34. D. Richman et al., Human immunodeficiency virus type 1 mutants resistant to nonnucleoside inhibitors of reverse transcriptase arise in tissue culture. Proc. Natl. Acad. Sci. U.S.A. 88, 11241-11245 (1991).

35. J. P. Barnard, K. D. Huber, N. Sluis-Cremer, Nonnucleoside Reverse Transcriptase Inhibitor Hypersusceptibility and Resistance by Mutation of Residue 181 in HIV-1 Reverse Transcriptase. Antimicrob. Agents Chemother. 63 (2019).

36. E. L. Asahchop et al., Distinct resistance patterns to etravirine and rilpivirine in viruses containing nonnucleoside reverse transcriptase inhibitor mutations at baseline. AIDS 27 (2013).

37. D. Richman et al., Human immunodeficiency virus type 1 mutants resistant to nonnucleoside inhibitors of reverse transcriptase arise in tissue culture. Proc. Natl. Acad. Sci. U.S.A. 88, 11241-11245 (1991).

38. V. Joly et al., Evolution of human immunodeficiency virus type 1 (HIV-1) resistance mutations in nonnucleoside reverse transcriptase inhibitors (NNRTIs) in HIV-1-infected patients switched to antiretroviral therapy without NNRTIs. Antimicrob. Agents Chemother. 48, 172-175 (2004).

39. M. A. Winters et al., Development of Elvitegravir Resistance and Linkage of Integrase Inhibitor Mutations with Protease and Reverse Transcriptase Resistance Mutations. PLoS One 7, e40514 (2012).

40. J. A. Fulcher, Y. Du, T. H. Zhang, R. Sun, R. J. Landovitz, Emergence of Integrase Resistance Mutations During Initial Therapy Containing Dolutegravir. Clin. Infect. Dis. 67, 791-794 (2018).

41. S. Fransen, S. Gupta, A. Frantzell, C. J. Petropoulos, W. Huang, Substitutions at Amino Acid Positions 143, 148, and 155 of HIV-1 Integrase Define Distinct Genetic Barriers to Raltegravir Resistance. J. Virol. 86, 7249-7255 (2012).

42. A. Fun et al., Impact of the HIV-1 genetic background and HIV-1 population size on the evolution of raltegravir resistance. Retrovirology 15, 1 (2018).

43. S. Y. Rhee et al., A systematic review of the genetic mechanisms of dolutegravir resistance. J Antimicrob Chemother 74, 3135-3149 (2019).

44. O. Goethals et al., Resistance Mutations in Human Immunodeficiency Virus Type 1 Integrase Selected with Elvitegravir Confer Reduced Susceptibility to a Wide Range of Integrase Inhibitors. J. Virol. 82, 10366-10374 (2008).

45. F. Canducci et al., Genotypic/phenotypic patterns of HIV-1 integrase resistance to raltegravir. J. Antimicrob. Chemother. 65, 425-433 (2010).

46. S. Gudipati, I. Brar, A. Golembieski, Z. Hanna, N. Markowitz, Occurrence of the S230R integrase strand inhibitor mutation in a treatment-naive individual case report. Medicine (Baltimore) 99, e20915 (2020).

47. L. K. Naeger, P. Harrington, T. Komatsu, D. Deming, Effect of Dolutegravir Functional Monotherapy on HIV-1 Virological Response in Integrase Strand Transfer Inhibitor Resistant Patients. Antivir. Ther. 21, 481-488 (2015).

48. C. Charpentier et al., Drug resistance profiles for the HIV integrase gene in patients failing raltegravir salvage therapy. HIV Med. 9, 765-770 (2008).

49. D. A. Cooper et al., Subgroup and Resistance Analyses of Raltegravir for Resistant HIV-1 Infection. N. Engl. J. Med. 359, 355-365 (2008).

50. J.-L. Blanco, V. Varghese, S.-Y. Rhee, J. M. Gatell, R. W. Shafer, HIV-1 Integrase Inhibitor Resistance and Its Clinical Implications. J. Infect. Dis. 203, 1204-1214 (2011).

51. M. Parczewski, D. Bander, A. Urbańska, A. Boroń-Kaczmarska, HIV-1 integrase resistance among antiretroviral treatment naive and experienced patients from Northwestern Poland. BMC Infect. Dis. 12, 368 (2012).

52. J. M. George et al., Rapid Development of High-Level Resistance to Dolutegravir With Emergence of T97A Mutation in 2 Treatment-Experienced Individuals With Baseline Partial Sensitivity to Dolutegravir. Open Forum Infectious Diseases 5, ofy221 (2018).

53. B. Atkinson, J. Isaacson, M. Knowles, E. Mazabel, A. K. Patick, Correlation between Human Immunodeficiency Virus Genotypic Resistance and Virologic Response in Patients Receiving Nelfinavir Monotherapy or Nelfinavir with Lamivudine and Zidovudine. J. Infect. Dis. 182, 420-427 (2000).

54. Y. Mitsuya et al., N88D Facilitates the Co-occurrence of D30N and L90M and the Development of Multidrug Resistance in HIV Type 1 Protease following Nelfinavir Treatment Failure. AIDS Res. Hum. Retroviruses 22, 1300-1305 (2006).

55. W. Sugiura et al., Interference between D30N and L90M in selection and development of protease inhibitor-resistant human immunodeficiency virus type 1. Antimicrob. Agents Chemother. 46, 708-715 (2002).

56. T. D. Wu et al., Mutation patterns and structural correlates in human immunodeficiency virus type 1 protease following different protease inhibitor treatments. J. Virol. 77, 4836-4847 (2003).

57. V. Varghese et al., Prototypical Recombinant Multi-Protease-Inhibitor-Resistant Infectious Molecular Clones of Human Immunodeficiency Virus Type 1. Antimicrob. Agents Chemother. 57, 4290-4299 (2013).

58. S. Y. Rhee, T. Liu, J. Ravela, M. J. Gonzales, R. W. Shafer, Distribution of human immunodeficiency virus type 1 protease and reverse transcriptase mutation patterns in 4,183 persons undergoing genotypic resistance testing. Antimicrob. Agents Chemother. 48, 3122-3126 (2004).

59. A. De Luca et al., Frequency and Treatment-Related Predictors of Thymidine-Analogue Mutation Patterns in HIV-1 Isolates after Unsuccessful Antiretroviral Therapy. J. Infect. Dis. 193, 1219-1222 (2006).

60. N. Yahi, C. Tamalet, C. Tourres, N. Tivoli, J. Fantini, "Mutation L210W of HIV-1 reverse transcriptase in patients receiving combination therapy. Incidence, association with other mutations, and effects on the structure of mutated reverse transcriptase" in J. Biomed. Sci. (2000), vol. 7, pp. 507-513.

61. C. A. Boucher et al., High-level resistance to (-) enantiomeric 2’-deoxy-3’-thiacytidine in vitro is due to one amino acid substitution in the catalytic site of human immunodeficiency virus type 1 reverse transcriptase. Antimicrob. Agents Chemother. 37, 2231-2234 (1993).

62. M. Tisdale, S. D. Kemp, N. R. Parry, B. A. Larder, Rapid in vitro selection of human immunodeficiency virus type 1 resistant to 3’-thiacytidine inhibitors due to a mutation in the YMDD region of reverse transcriptase. Proc. Natl. Acad. Sci. U.S.A. 90, 5653-5656 (1993).

63. W. Keulen, C. Boucher, B. Berkhout, Nucleotide substitution patterns can predict the requirements for drug-resistance of HIV-1 proteins. Antiviral Res. 31, 45-57 (1996).

64. S. D. Frost, M. Nijhuis, R. Schuurman, C. A. Boucher, A. J. Brown, Evolution of lamivudine resistance in human immunodeficiency virus type 1-infected individuals: the relative roles of drift and selection. J. Virol. 74, 6262-6268 (2000).

65. R. W. Shafer et al., Combination therapy with zidovudine and didanosine selects for drug-resistant human immunodeficiency virus type 1 strains with unique patterns of pol gene mutations. J. Infect. Dis. 169, 722-729 (1994).

66. L. R. Miranda, M. Gotte, F. Liang, D. R. Kuritzkes, The L74V mutation in human immunodeficiency virus type 1 reverse transcriptase counteracts enhanced excision of zidovudine monophosphate associated with thymidine analog resistance mutations. Antimicrob. Agents Chemother. 49, 2648-2656 (2005).

67. V. Trivedi et al., Impact of human immunodeficiency virus type 1 reverse transcriptase inhibitor drug resistance mutation interactions on phenotypic susceptibility. AIDS Res. Hum. Retroviruses 24, 1291-1300 (2008).

68. G. L. Melikian et al., Non-nucleoside reverse transcriptase inhibitor (NNRTI) cross-resistance: implications for preclinical evaluation of novel NNRTIs and clinical genotypic resistance testing. J. Antimicrob. Chemother. 69, 12-20 (2014).

69. N. T. Parkin, S. Gupta, C. Chappey, C. J. Petropoulos, The K101P and K103R/V179D mutations in human immunodeficiency virus type 1 reverse transcriptase confer resistance to nonnucleoside reverse transcriptase inhibitors. Antimicrob. Agents Chemother. 50, 351-354 (2006).

70. E. Magiorkinis et al., Emergence of an NNRTI resistance mutation Y181C in an HIV-infected NNRTI-naive patient. AIDS Res. Hum. Retroviruses 24, 413-415 (2008).

71. H. T. Xu et al., Molecular mechanism of antagonism between the Y181C and E138K mutations in HIV-1 reverse transcriptase. J. Virol. 86, 12983-12990 (2012).

72. W. Guo et al., Impact of Y181C and/or H221Y mutation patterns of HIV-1 reverse transcriptase on phenotypic resistance to available non-nucleoside and nucleoside inhibitors in China. BMC Infect. Dis. 14, 237 (2014).

73. T. Shirasaka et al., Emergence of human immunodeficiency virus type 1 variants with resistance to multiple dideoxynucleosides in patients receiving therapy with dideoxynucleosides. Proc. Natl. Acad. Sci. U.S.A. 92, 2398-2402 (1995).

74. R. W. Shafer et al., Drug resistance and heterogeneous long-term virologic responses of human immunodeficiency virus type 1-infected subjects to zidovudine and didanosine combination therapy. The AIDS Clinical Trials Group 143 Virology Team. J. Infect. Dis. 172, 70-78 (1995).

75. A. K. Iversen et al., Multidrug-resistant human immunodeficiency virus type 1 strains resulting from combination antiretroviral therapy. J. Virol. 70, 1086-1090 (1996).

76. K. Das, S. E. Martinez, E. Arnold, Structural Insights into HIV Reverse Transcriptase Mutations Q151M and Q151M Complex That Confer Multinucleoside Drug Resistance. Antimicrob Agents Chemother 61 (2017).

77. P. L. Tzou et al., Integrase strand transfer inhibitor (INSTI)-resistance mutations for the surveillance of transmitted HIV-1 drug resistance. J. Antimicrob. Chemother. 75, 170-182 (2020).

78. N. A. Margot, R. R. Ram, K. L. White, M. E. Abram, C. Callebaut, Antiviral activity of HIV-1 integrase strand-transfer inhibitors against mutants with integrase resistance-associated mutations and their frequency in treatment-naive individuals. J. Med. Virol. 91, 2188-2194 (2019).

79. N. J. Cook et al., Structural basis of second-generation HIV integrase inhibitor action and viral resistance. Science 367, 806-810 (2020).

80. M. Li et al., Mechanisms of HIV-1 integrase resistance to dolutegravir and potent inhibition of drug-resistant variants. Sci. Adv. 9, eadg5953 (2023).

81. S. Reigadas et al., The HIV-1 integrase mutations Y143C/R are an alternative pathway for resistance to Raltegravir and impact the enzyme functions. PLoS One 5, e10311 (2010).

82. Z. Hu, D. R. Kuritzkes, Effect of raltegravir resistance mutations in HIV-1 integrase on viral fitness. J. Acquir. Immune. Defic. Syndr. 55, 148-155 (2010).

